# cuBNM: GPU-Accelerated Brain Network Modeling

**DOI:** 10.1101/2025.11.13.688224

**Authors:** Amin Saberi, Bin Wan, Kevin J. Wischnewski, Kyesam Jung, Leonard Sasse, Felix Hoffstaedter, Boris C. Bernhardt, Simon B. Eickhoff, Oleksandr V. Popovych, Sofie L. Valk

## Abstract

Brain network modeling uses computer simulations to infer about latent neural properties at micro- and mesoscales by fitting brain dynamic models to empirical data of individual subjects or groups. However, computational costs of (individualized) model fitting is a major bottleneck, limiting the practical feasibility of this approach to larger cohorts and more complex models, and highlighting the need for scalable simulation implementations. Here, we introduce *cuBNM*, a Python package which leverages parallel processing of graphics processing units to massively accelerate simulations of brain network models. We show running simulations on graphics processing units is several hundred times faster compared to central processing units. We demonstrate the usage of cuBNM by running optimization of group-level and individualized low- and high-dimensional models. As examples of the utility of individualized models, we investigated test-retest reliability and heritability of simulated and empirical measures in the Human Connectome Project dataset. We found simulated features were fairly reliable and significantly heritable, suggesting their biological plausibility. Overall, cuBNM enables large-scale simulations of brain network models, opening new avenues for studying latent neural processes across diverse populations, dense networks, and high-dimensional models, which was previously impractical due to computational constraints.

## 1 Introduction

Understanding the neural mechanisms underlying human cognition and behavior, and their alterations in disorders, requires studying the brain across multiple spatial and temporal scales^1^. Yet, fundamental processes at the micro- and mesoscales, such as microcircuit dynamics^2^, excitation-inhibition balance^3^, and neural gain^4^, cannot be readily measured *in vivo* in humans due to ethical and technical constraints. In contrast, noninvasive neuroimaging techniques provide scalable and accessible *in vivo* measures of brain function and dynamics, though at the macroscale. Therefore, there is an explanatory gap between the observed brain function at macroscale and the underlying neural mechanisms of interest at the micro- and mesoscales. This highlights the need for experimental^5^ and theoretical neuroscience approaches^2,6,7^ that can bridge these scales of investigation.

Brain network modeling (BNM)^a^ provides a computational framework to investigate how macroscale dynamics of the brain emerge from mesoscale activity and interactions of neural populations across regions^2,6,7^. In this approach, local neural models are coupled within a structural scaffold and simulated to generate observation-level signals such as blood-oxygen-level-dependent (BOLD) signals or magnetoencephalography/electroencephalography recordings. By fitting BNMs to empirical data of individuals or groups, representative models can be constructed. These models can offer *in silico* insights into latent neural properties at micro-/mesoscales, such as excitation-inhibition balance^8–10^, or neural gain^11,12^. These latent properties can then be examined in relation to behavioral phenotypes^13^, developmental trajectories^8,9^, or neuropsychiatric disorders^10,14^.

A critical challenge in this framework is scalable model fitting, i.e., tuning of model parameters to generate simulated data that most closely resembles the empirical data of a large number of subjects or groups. Model fitting often requires evaluating several thousands of parameter combinations per subject, to adequately explore the parameter space. Each evaluation involves running a simulation, i.e., numerical integration of coupled stochastic differential equations that define the BNM. Consequently, fitting BNMs to empirical data is computationally costly, creating a major bottleneck for their application to larger cohorts, denser networks, or more complex models. As a result, researchers have often needed to make practical trade-offs, such as limiting the number of subjects, using coarser atlases, or simplifying model complexity, to maintain feasibility within available computing resources. This challenge highlights the need for more efficient BNM simulation approaches.

Traditionally, BNM simulations have been primarily implemented on central processing units (CPUs)^15,16^. However, CPU-based implementations are typically constrained by relatively limited parallel processing capacity, as simulations, nodes, and time steps are processed sequentially and can only be parallelized across simulations to the extent of the number of available CPU threads. This constraint poses a practical challenge to scalability of BNMs and limits the feasibility and cost-efficiency of large-scale model optimizations. Graphics processing units (GPUs), in contrast, are inherently designed for parallel processing and can provide a more efficient solution. In GPU-based implementation, calculations across simulations and network nodes can be performed concurrently (Fig. 1). This can substantially reduce computation times, making large-scale BNM optimizations more practical and accessible. For instance, a typical study including a few hundred subjects, depending on the model complexity, network density, and simulation length, may require hundreds of thousands of CPU core-hours (with multi-threaded processing), but only a few thousand GPU-hours.

**Fig. 1.**
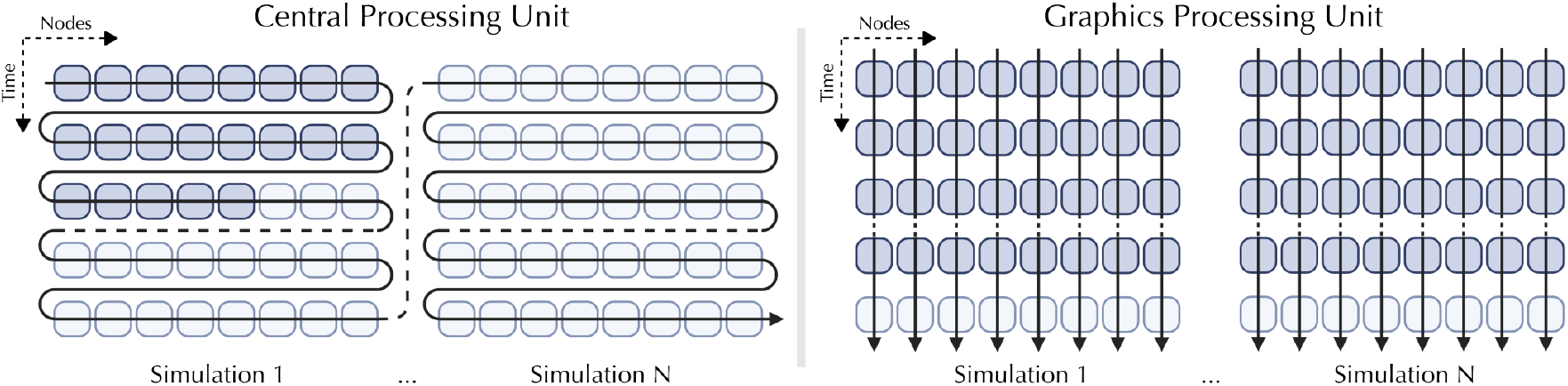
Schematic illustration of brain network model simulations on central processing units and graphics processing units. Each cell represents a network node at a given simulation time in a given simulation. Dark-colored cells indicate completed calculations, and arrows show the order of execution. Created in BioRender. Saberi, A. (2026).

**Fig. 2.**
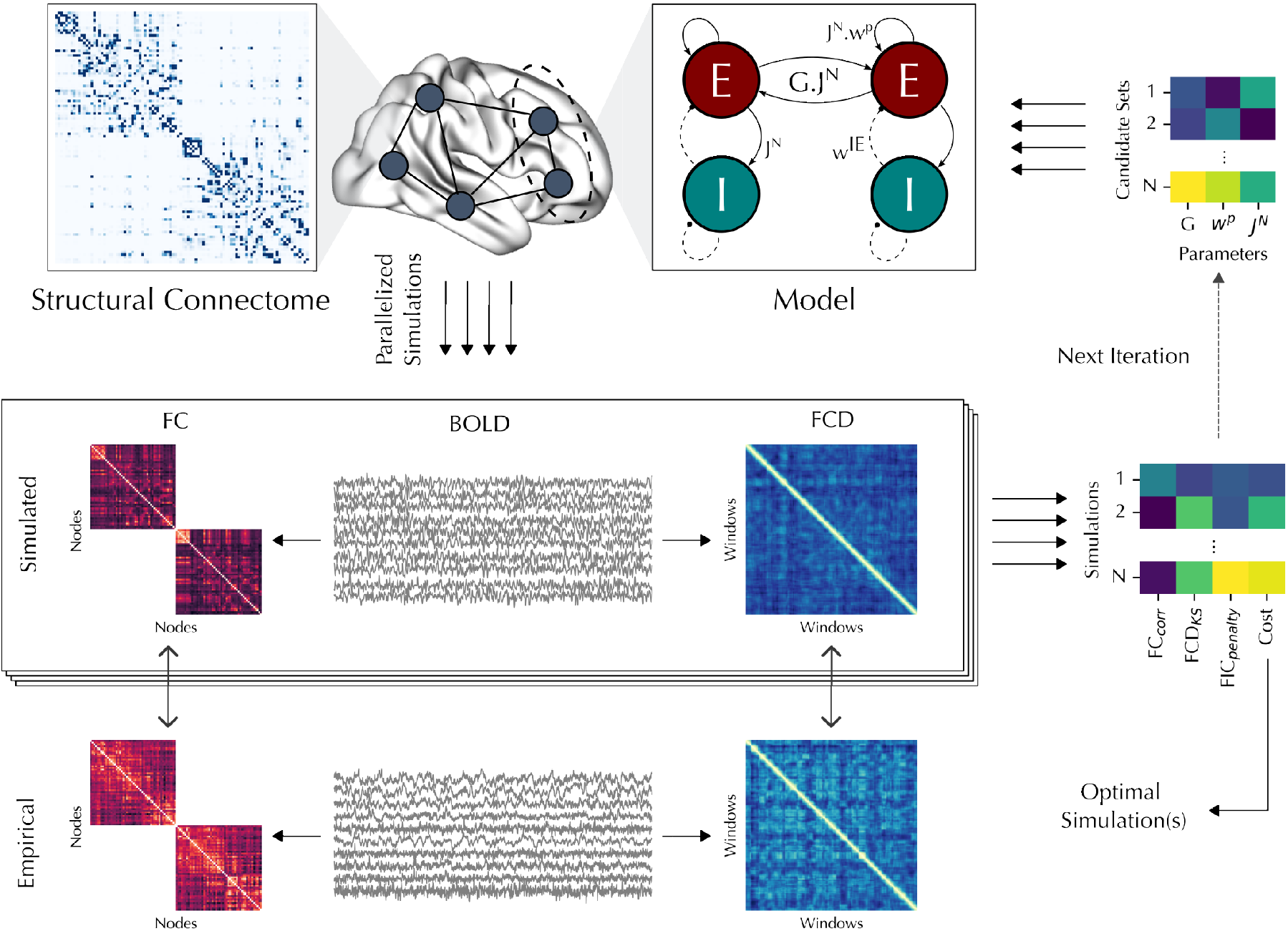
Optimization of brain network models. A brain network model describes the dynamics of a network of nodes connected via a structural connectome. As an example, the reduced Wong-Wang model is shown, with three free parameters: *G, w*^*p*^, and *J*^*N*^. The goal of optimization is to fit model parameters to empirical data, such as blood-oxygen-level-dependent (BOLD) signal. Candidate parameter sets are sampled, and parallel simulations are run for each set. The simulated BOLD signals are compared to empirical data using evaluation metrics such as the correlation between simulated and empirical functional connectivity (FC) matrices (*FC*_*corr*_) and the Kolmogorov-Smirnov distance between simulated and empirical functional connectivity dynamics (FCD) matrices (*FCD*_*KS*_). Additional penalty terms may also be included (e.g., the feedback inhibition control (FIC) penalty in the reduced Wong-Wang model). The evaluation metrics are combined to define the cost function of each simulation. In evolutionary optimization, new parameter sets are iteratively sampled based on cost functions from previous iterations, whereas in grid search, all parameter combinations are simulated in a single run on a predefined parameter grid. After completion of the optimization procedure, the optimal simulation is defined as the one with the lowest cost function.

Indeed, there has been a growing interest in using GPUs to accelerate numerical simulation of dynamical systems across domains^17–20^. Yet, while several GPU-based implementations of BNM simulations have been introduced^21–29^, the application of GPUs for scalable simulation-optimization of BNMs remains underexplored. Here, we introduce *cuBNM*, a Python package with a C++ and CUDA backend designed specifically for the computational requirements of BNMs, facilitating large-scale and individualized modeling by leveraging GPUs to accelerate BNM simulations and parameter optimization. cuBNM parallelizes calculations across subjects, simulations, and nodes, and extends GPU acceleration beyond numerical integration to the computation of derived features such as simulated FC and FCD and their fit to empirical data. It integrates grid search and evolutionary optimization, supports homogeneous and heterogeneous parameterization, and provides a Python application programming interface (API) and command-line interface (CLI). In this paper we provide tutorials and example applications to demonstrate its use. We show how GPU acceleration facilitates fitting low- or high-dimensional BNMs using both grid search and evolutionary optimization approaches. As examples of the utility of individualized models, we create BNMs for individuals from the Human Connectome Project - Young Adults (HCP-YA)^30,31^, compare model fit across homogeneous and heterogeneous parameterizations, and evaluate test-retest reliability and heritability of simulated and empirical measures, providing evidence for the biological relevance of individualized BNMs and illustrating analyses that scalable modeling makes feasible. Finally, we evaluate scaling of computational time as a function of number of simulations and network size, showing that the GPU-based implementation achieves speed-ups of up to 1,266x compared to the single-threaded CPU-based implementation, i.e., a single GPU having the performance of 1,266 CPU threads.

## 2 Results

cuBNM is an open-source Python package with a C++ and CUDA backend that enables efficient GPU-accelerated simulations and optimization of BNMs. In addition to GPUs, it supports execution of the simulations on CPUs via an efficient backend written in C++. The package can be installed via: `pip install cubnm`. It provides both Python API and CLI. Comprehensive documentation and tutorials are available at https://cubnm.readthedocs.io.

### 2.1 Defining a brain network modeling problem

A BNM consists of network nodes (brain areas) interconnected through a structural connectome (SC). Nodes are typically defined based on anatomical^32^, functional^33^, or multimodal^34^ atlases at varying levels of granularity. SC quantifies the pattern and strength of white matter (and sometimes, lateral) connections between the nodes, and is typically derived from diffusion-weighted imaging (DWI), based on the data of an individual or a group of subjects.

BNMs mathematically describe the activity of each node, including its intrinsic and often stochastic activity, in interaction with the other nodes in the network. cuBNM includes a number of predefined BNMs (Table S1) and supports custom BNM implementations. This paper focuses on the reduced Wong-Wang (rWW) BNM with feedback inhibition control (FIC)^35,36^. The rWW model consists of interconnected excitatory (E) and inhibitory (I) neuronal ensembles and is controlled by free parameters *G* (global coupling), 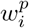 (local excitatory recurrence of node *i*), and 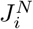 (N-methyl-D-aspartate [NMDA] receptor conductance of node *i*) (Equations 1–6, Table S2).

Brain network modeling can be used to explore how parameter variations influence network dynamics *in silico*, or to optimize parameters to best match empirical functional data, such as resting-state functional magnetic resonance imaging (rs-fMRI), in order to construct models representing a group or subject. To compare with empirical data, a forward model such as the Balloon-Windkessel model^37^ transforms simulated neuronal activity into observation-level (e.g. BOLD) signals. Goodness-of-fit (GOF) is then quantified using metrics such as functional connectivity (FC) correlation (*FC*_*corr*_) and functional connectivity dynamics (FCD) distance (*FCD*_*KS*_), which combined with optional penalty terms (e.g., the FIC penalty in the rWW model) define the cost function of a given simulation (Equation 14). In model fitting, the free parameters are tuned aiming to minimize the cost function.

In cuBNM, model components, simulation configurations, and optimization targets can be specified as a “BNM problem” using the `BNMProblem` class (Section 4.4). The BNM problem is passed to optimization algorithms that use GPU acceleration to efficiently search for optimal parameter sets that minimize the cost function. In the following sections, we will illustrate the utility of this framework by using different optimization and model parameterization approaches to fit the rWW BNM to the group-averaged as well as individual-level data of subjects from the HCP-YA dataset. In all models, the network consisted of 100 nodes defined based on the Schaefer-100 atlas^33^ and simulations were run for a duration of 900 s (BOLD repetition time [TR] = 0.72 s, integration step = 0.1 ms).

### 2.2 Grid search and evolutionary optimization of BNMs

Fitting BNMs to empirical data in cuBNM can be performed using two optimization approaches: grid search and evolutionary optimization. Grid search exhaustively evaluates all parameter combinations across a predefined range and resolution, whereas evolutionary optimizers selectively sample the parameter space across generations, iteratively evolving towards an optimal solution. In cuBNM, the full grid in grid search and each generation of evolutionary optimization are parallelized to the extent of hardware capacity. Here, to demonstrate the usage of these optimization algorithms in cuBNM and to introduce their underlying concepts, we applied both approaches to a homogeneous low-dimensional rWW model with three free parameters *G, w*^*p*^, and *J*^*N*^. Importantly, in this homogeneous model, unlike the heterogeneous models covered in the next section, the regional parameters *w*^*p*^ and *J*^*N*^ were assumed to be constant across nodes. The model was fit to the group-averaged data of HCP-YA training sample (706 subjects; age: 28.7*±*3.7; 54.1% female), and performance was additionally evaluated on the independent test sample (303 subjects; age: 28.8*±*3.7; 54.1% female).

In the grid search (Fig. 3), we sampled each parameter at 22 evenly spaced values, running a total of 22^3^ = 10,648 simulations. Running the full grid search took 32.2 minutes on an Nvidia A100 GPU (40 GB-SXM), whereas a single simulation on a CPU core required 161 s, projecting it to take 19.5 core-days for the full grid, indicating a ∼872-fold GPU speed-up. This means that a single GPU was ∼872 times faster than a single CPU thread, or in other words, a single GPU had the performance of ∼872 CPU threads (see subsection 2.5 for more comprehensive scaling analyses including comparisons with multi-core CPU). Notably, we verified in an independent smaller grid search (running 1000 simulations with durations of 60 s) that GPU and CPU implementations result in identical outputs (Supplementary Text A.1).

**Fig. 3.**
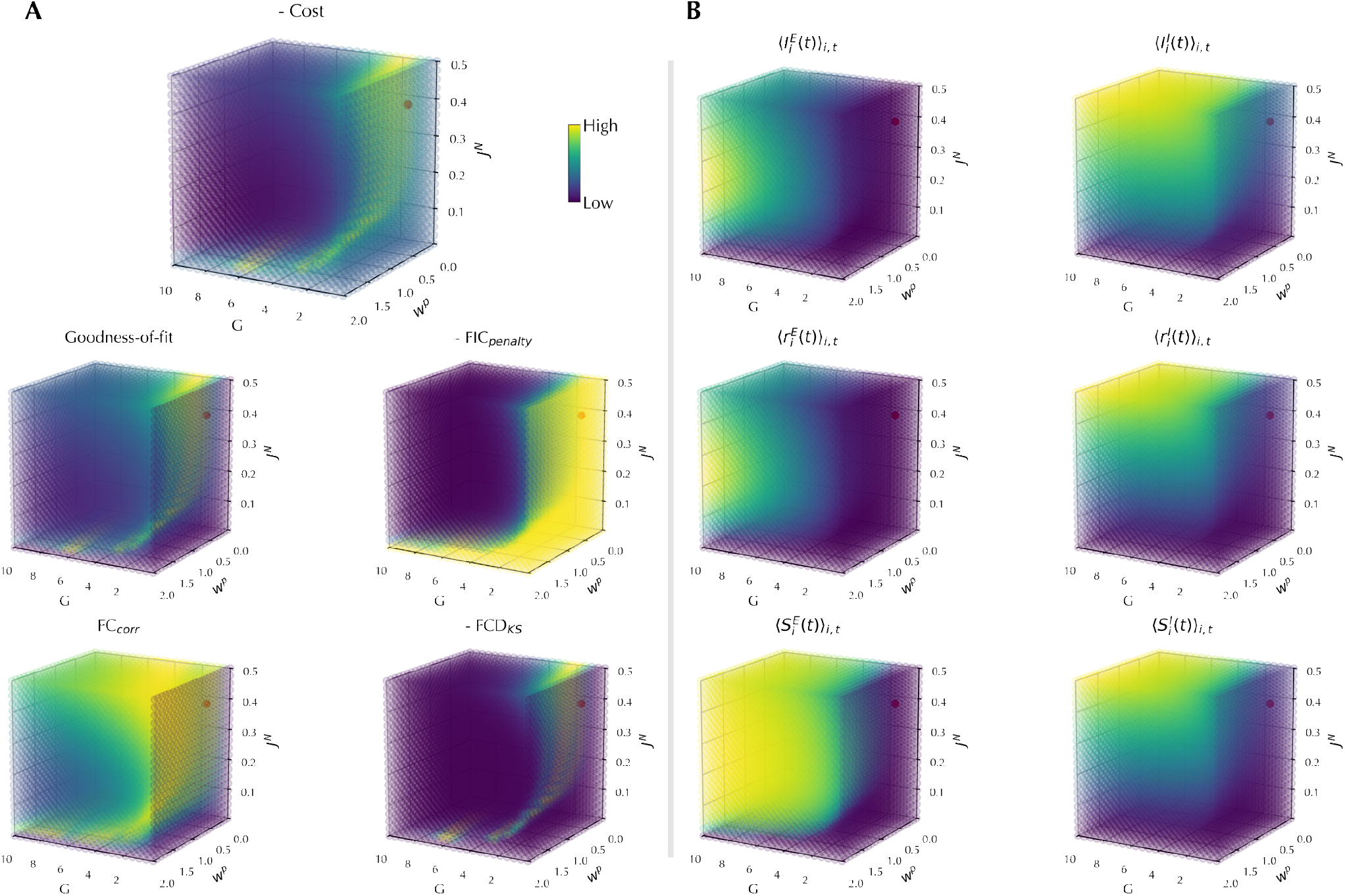
Grid search of the homogeneous model. Three-dimensional grids show the variation of cost function and its components (**A**) as well as the average state variables (**B**) as a function of the parameters *G, w*^*p*^, and *J*^*N*^. Each dimension is sampled at 22 evenly spaced points. Red spheres mark the optimum, i.e., the simulation with the lowest cost. In **A**, the cost function, *FIC*_*penalty*_ and *FCD*_*KS*_ are negated so that in all grids brighter colors indicate better fits.

The global optimum was identified at *G* = 1.429, *w*^*p*^ = 0.095, and *J*^*N*^ = 0.381, with a cost of -0.119 on the training data and -0.118 on the test data (Fig. S1, Table S3). Beyond the optimal point, grid search provides a thorough overview of the parameter space. This makes it useful for studying how model behavior, including the model fit to the data (Fig. 3A), as well as its state variables (Fig. 3B), changes across parameters. By visualizing the grid of cost function and its components we can investigate the relationship between the parameters, their potential redundancies, and locations and shapes of local optimal areas. For example, in this grid we observe a locally optimal area as curved sheet near an edge of the parameter space with lower *G* values, and another smaller locally optimal region with high *G* and *w*^*p*^ but lower *J*^*N*^ values (Fig. 3A). In addition, visualizing the grids of average state variables shows, for example, that the mean excitatory input current 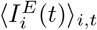 increased with *G* and *w*^*p*^ but showed a U-shaped relationship with *J*^*N*^ (Fig. 3B, Fig. S2), likely due to the dual excitatory and inhibitory action of NMDA conductance in the rWW model.

We next applied an evolutionary optimization approach to the same BNM problem (Fig. 4). cuBNM supports a broad set of optimizers from the *PyMoo* library^38^. In this paper we focus on the covariance matrix adaptation evolution strategy (CMA-ES)^39^. Here, we applied CMA-ES using 128 particles per generation with a maximum of 120 generations, subject to early termination criteria. The optimizer terminated early after 27 generations (Fig. 4), running a total of 3,457 simulations (GPU time: 20.0 minutes; projected CPU time: 6.3 core-days; GPU speed-up: 455x). CMA-ES identified an optimal point at *G* = 1.540, *w*^*p*^ = 0.006, and *J*^*N*^ = 0.408, yielding a cost of -0.125 on the training data and -0.124 on the test data (Fig. S1, Table S3), slightly better than the grid search optimum.

**Fig. 4.**
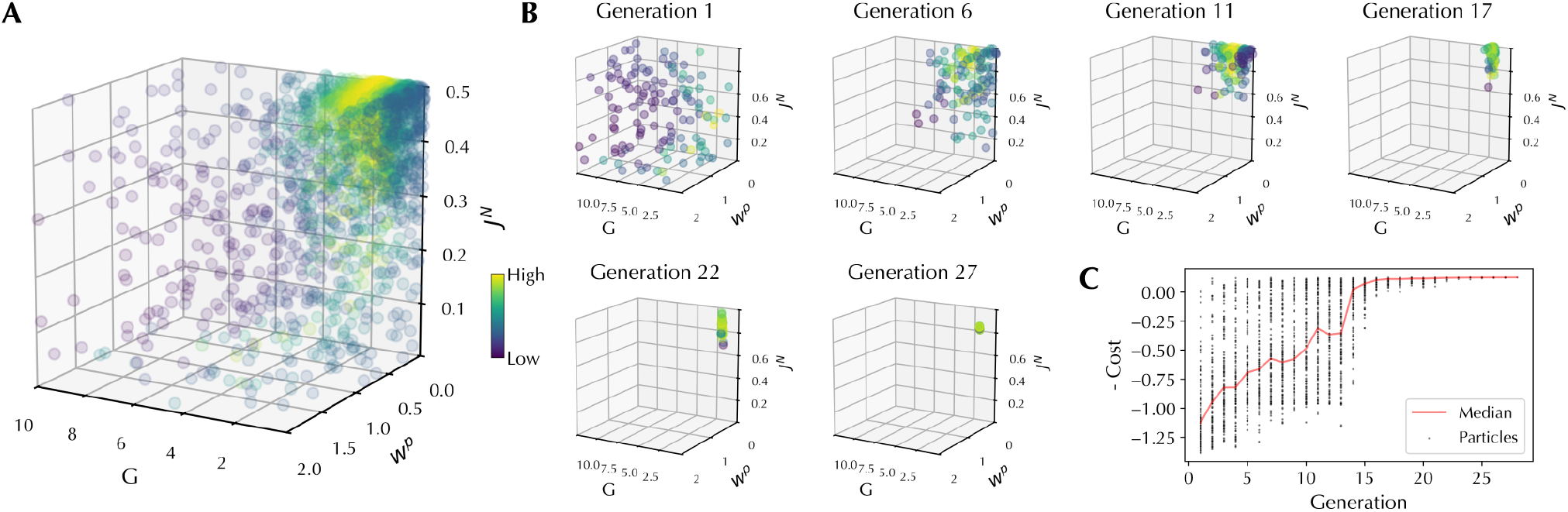
Evolutionary optimization of the homogeneous model. (**A**) Three-dimensional visualization of negative cost values across all parameter combinations evaluated during CMA-ES optimization. (**B**) Three-dimensional visualization of parameter combinations sampled in eight selected generations, showing gradual convergence of the optimizer toward the optimal region of the parameter space. (**C**) Evolution of negative cost across particles (dots) and generations. The red line indicates the median negative cost per generation.

Overall, the choice of grid search or evolutionary approaches for fitting BNMs depends on the study goal and dimensionality of the model. Grid search is particularly useful when a comprehensive overview of the parameter space is of interest, enabling the study of parameter interactions, redundancies, and the landscape of local optima. However, its brute-force nature makes it computationally infeasible for higher-dimensional models due to the curse of dimensionality, and its resolution is inherently limited by the discretization of the parameter space. In contrast, evolutionary optimization is more efficient when the primary aim is to identify global optimal points, and is in practice the only feasible option for higher-dimensional models.

### 2.3 High-dimensional models with heterogeneous regional parameters

The homogeneous model used in the previous section assumes that within each simulation the local parameters *w*^*p*^ and *J*^*N*^ are identical across all nodes. This assumption is biologically unrealistic, as previous research has shown microcircuit and synaptic features vary across areas^12,40,41^. We can account for this heterogeneity in the model by allowing nodes within the same simulation to have different regional parameter values. This can be done in two ways: (i) **Map-based heterogeneity**: Regional parameters varying systematically constrained by one or more fixed maps^8,9,12,36,42^ (Fig. 5A), or (ii) **Node-based heterogeneity**: Regional parameters varying independently across nodes or node groups^10,43,44^ (Fig. 5B). Notably, both approaches inevitably increase the complexity of the model and the number of free parameters. While low-dimensional models can be fit using both grid search and evolutionary optimizers, grid search quickly becomes infeasible as dimensionality increases. In such cases, it is crucial to efficiently and selectively explore the parameter space with evolutionary optimization. To illustrate these approaches, here, we fit the two types of heterogeneous rWW BNMs to the group-averaged HCP-YA training data using the CMA-ES optimization algorithm with 128 particles per generation and a maximum of 120 generations, subject to early termination criteria.

**Fig. 5.**
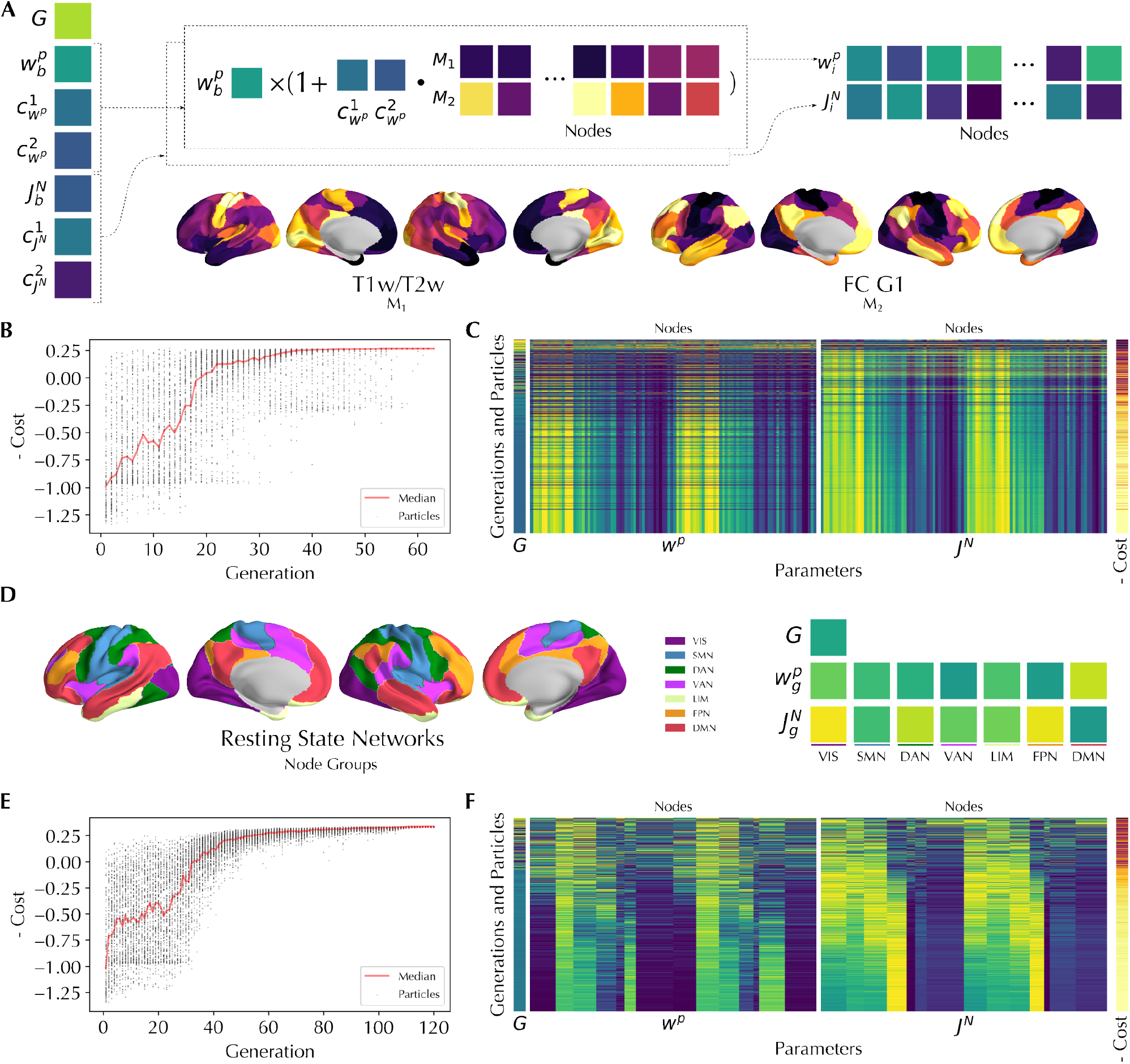
Evolutionary optimization of heterogeneous models. (**A**) In the map-based heterogeneous model, regional parameters 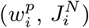 are determined based on a parametric combination of spatial maps of heterogeneity (e.g., T1w/T2w ratio, principal gradient of FC) using a set of bias terms (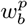 _and 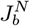_) and coefficients (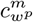 and 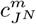 for each map *m*). (**D**) In the node-based heterogeneous model, nodes are grouped based on a categorical map (e.g., the canonical resting-state networks), with separate free parameters 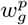 and 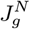 for each group. (**B, E**) Evolution of negative cost values across CMA-ES particles (dots) and generations in map-based (**B**) and node-based (**E**) heterogeneous models. Red line indicates the median per generation. (**C, F**) Evolution of global and regional parameter values with their associated cost functions across all sampled particles arranged from early (top) to late (bottom) generations, showing the gradual convergence of the optimizer, in map-based (**C**) and node-based (**F**) heterogeneous models

#### 2.3.1 Map-based heterogeneous model

In this model the regional parameters were defined according to two biological, fixed maps, previously calculated using the HCP-YA dataset: (i) the T1w/T2w ratio map and (ii) the first principal gradient of FC (FC G1). These maps represent cortical microstructural and functional variability and have been commonly used in BNM studies to introduce heterogeneity of regional parameters^8,9,12,36,42^. A parametric combination of the maps, based on a set of bias terms (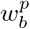 and 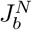) and coefficients (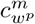 and 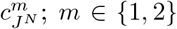) determines the value of regional parameters 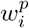 and 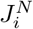 in each node *i* (Fig. 5A). Consequently, these bias terms and coefficients, together with global coupling *G*, are the 7 tunable free parameters of the model. The CMA-ES optimization on this model stopped early after 62 generations (running a total of 7,937 simulations within 42.6 minutes on GPU; projected CPU time: 14.5 core-days; GPU speed-up: 492x; Fig. 5B, C), leading to an optimal simulation with a cost of -0.267 on the training data -0.258 on the test data (Fig. S1, Table S3).

#### 2.3.2 Node-based heterogeneous model

Here, we grouped the nodes into seven groups based on the canonical resting-state network that they belong to^45^. Each group g ∈ {1, …, 7} was assigned its own pair of regional parameters, 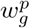 and 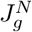, which were shared across all nodes within that network (Fig. 5D). Thereby, together with the global coupling *G*, this model has 15 free parameters. The CMA-ES optimization on this model continued for 120 generations (running a total of 15,360 simulations within 144.4 minutes on GPU; projected CPU time: 28.2 core-days; GPU speed-up: 480x; Fig. 5E, F), leading to an optimal simulation with a cost of -0.338 on the training data and -0.327 on the test data (Fig. S1, Table S3).

Overall, in both heterogeneous models, compared to the homogeneous model, we observed a considerable increase in the model fit to the empirical data.

### 2.4 Building individualized BNMs

Thus far, we illustrated the functionality of cuBNM, by fitting the homogeneous and heterogeneous BNMs to group-averaged data using grid search or CMA-ES. While such group-level models are useful, many research questions concern intra- or inter-individual variability of simulation-derived features. Addressing these questions requires individualized models fitted to subject- or session-specific data. The GPU-accelerated implementation of BNM simulations in cuBNM is particularly suited for this purpose, enabling the scaling of BNM framework to individuals across entire cohorts. Indeed, this highlights a key advantage of cuBNM, making it a practical and accessible tool for investigation of biologically relevant intra- and inter-individual differences in latent neural features, such as excitation–inhibition ratio^8^.

Here, to illustrate this functionality, we used the individual SCs and session-specific functional data of 430 twin subjects (age: 29.2*±*3.3; 59.1% female) from the HCP-YA dataset and constructed subject- and session-specific BNMs. As illustrative examples of the utility of individualized models, we aimed to address three questions using this data: (i) How does the fit of optimal simulations differ between homogeneous, map-based heterogeneous, and node-based heterogeneous models, at the level of individuals? (ii) What is the test-retest reliability of empirical and simulated features across two sessions? (iii) To what extent are simulation-derived features heritable?

To reduce convergence to local optima, for each model, optimization was performed twice using different optimizer random seeds, and the better of the two optima was selected. In total, 2,074 CMA-ES optimization runs were performed, including 28,158,401 simulations. These runs were completed in 1,426.0 hours on Nvidia A100 GPUs (average 182 ms per simulation), compared to an estimated 141.4 CPU core-years, corresponding to a GPU speed-up of 869x. Optimizations were executed in batches of 16 parallel runs (2,048 parallel simulations) per GPU to maximize parallelization.

#### 2.4.1 Comparison of homogeneous and heterogeneous models across individuals

We earlier observed incremental improvements in model fit from homogeneous to map-based and node-based heterogeneous models when using group-averaged models. To test this at the individual level, we fit the three model types to day 1 data of a set of 96 unrelated subjects (age: 29.2*±*3.2; 60.4% female). Paired t-tests showed an improvement in model fit with increased complexity of the model, with node-based heterogeneous models performing best (cost = -0.271*±*0.062, *FC*_*corr*_ = 0.433*±*0.066, *FCD*_*KS*_ = 0.142*±*0.038, *FIC*_*penalty*_ = 0.019*±*0.007), followed by map-based heterogeneous models (cost = -0.193*±*0.074, *FC*_*corr*_ = 0.376*±*0.082, *FCD*_*KS*_ = 0.159*±*0.040, *FIC*_*penalty*_ = 0.024*±*0.023), and homogeneous models performing worst (cost = -0.052*±*0.102, *FC*_*corr*_ = 0.225*±*0.069, *FCD*_*KS*_ = 0.141*±*0.062, *FIC*_*penalty*_ = 0.032*±*0.016) (Fig. 6A). This mirrors the group-level findings. Given the better performance of the node-based heterogeneous model, we used this model in the subsequent test-retest reliability and heritability analyses.

**Fig. 6.**
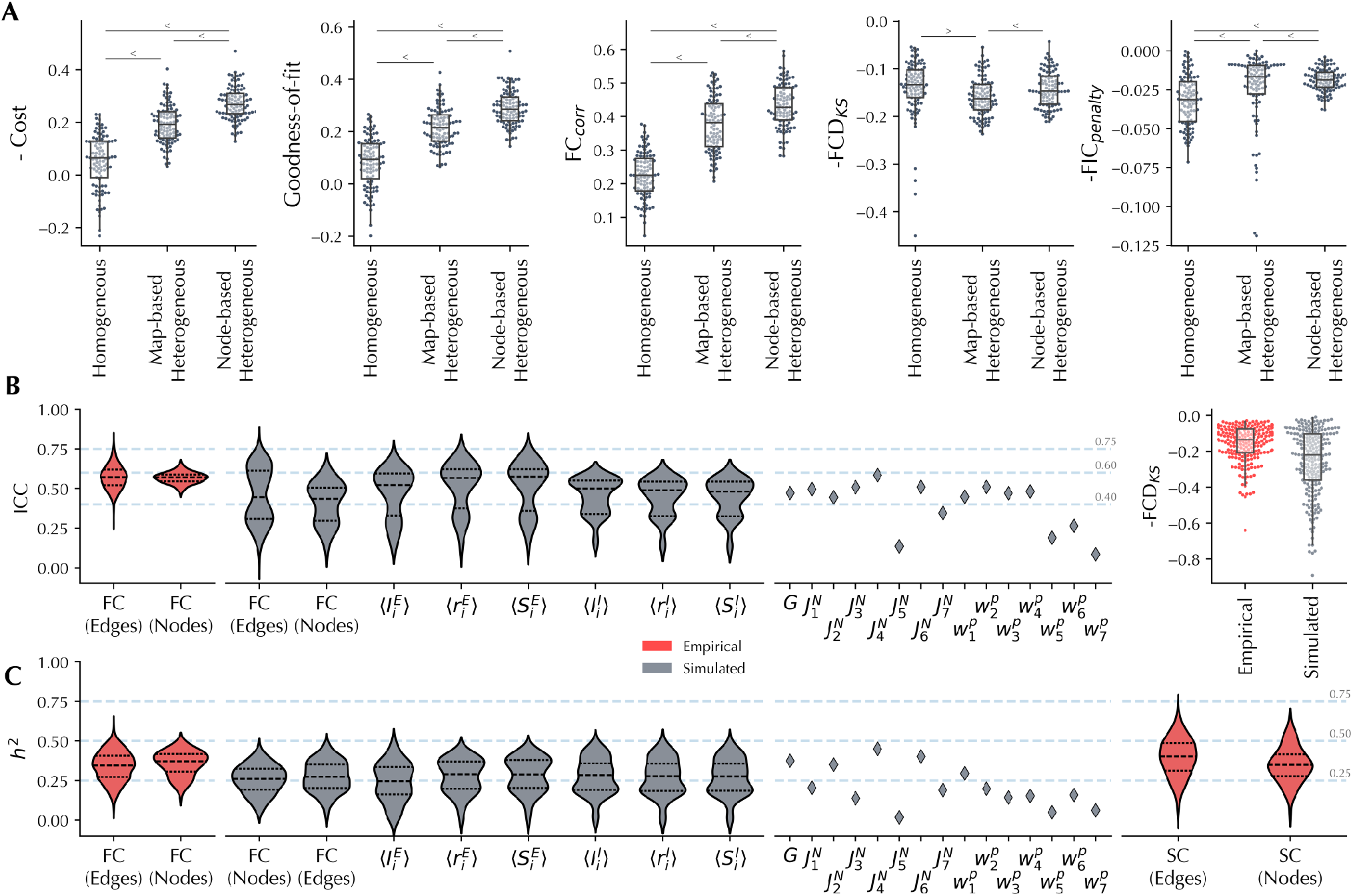
Model fit, reliability, and heritability of individualized BNMs. (**A**) Comparison of negative cost and its components across homogeneous, map-based heterogeneous, and node-based heterogeneous models. Dots denote the optimal simulations of each subject and model. Horizontal lines indicate significant pairwise differences between models, with the arrow direction showing the better fit. (**B**) *Left* : Intraclass correlation coefficients (ICCs) for empirical (red) and simulated (grey) features across day 1 and day 2 sessions. *Right* : Kolmogorov-Smirnov (KS) distance of functional connectivity dynamics (FCD) distributions between sessions for empirical (red) and simulated (grey) data. (**C**) Heritability (*h*^2^) of empirical (red) and simulated (grey) features based on the averaged data from both sessions. In **B** and **C**, violin plots represent the distribution of values across connectome edges or network nodes, with black lines indicating medians and quartiles. Diamonds indicate point estimates.

#### 2.4.2 Test-retest reliability of empirical and simulated features

We next quantified the test-retest reliability of empirical and simulated features across the two imaging sessions (days 1 and 2) using intraclass correlation coefficients (ICCs) (Fig. 6B). In this analysis, we used the data from 217 unrelated subjects (age: 29.3*±*3.4; 59.4% female).

Empirical FC showed fair average reliability both at the level of edges (0.570*±*0.074) and node-wise strengths (0.567*±*0.035). The average reliability of simulated FC was also in the fair range (edge-level: 0.455*±*0.173; node-level: 0.409*±*0.128), but it was significantly lower than that of empirical FC (edge-level: mean difference = -0.115, T = -33.48, p < 0.001; node-level: mean difference = -0.158, T = -12.40, p < 0.001). Simulated time-averaged state variables showed fair average reliability 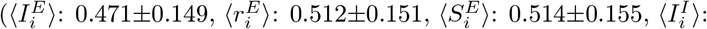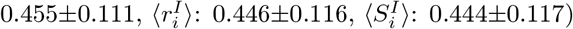. Notably, we observed patterns of regional variability of simulated FC and excitatory state variables were significantly (p_*spin,F DR*_ < 0.05) co-aligned with the spatial pattern of node-wise SC-FC coupling, highlighting the influence of SC in the simulated outcomes (Fig. S3). Model parameters showed poor-to-fair reliability with *G* and 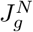 being generally more reliable than 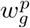. To evaluate cross-session stability of empirical and simulated FCDs, we compared Kolmogorov-Smirnov distances between session-specific distributions, and found reasonable stability in both empirical (*FCD*_*KS*_ = 0.157*±*0.102) and simulated (*FCD*_*KS*_ = 0.256*±*0.182) data, with empirical FCDs being more stable (mean difference = 0.099, T = -16.27, p < 0.001).

Together, these results indicate that many simulated features had fair reliability, but empirical features were more reliable.

#### 2.4.3 Heritability of empirical and simulated features

Following the assessment of test–retest reliability, we next sought to determine to what extent the simulated and empirical features are heritable. This analysis aimed to disentangle genetic and environmental contributions to inter-individual variability of these features. We estimated the heritability (*h*^2^) of empirical and simulated features based on the averaged data of the two sessions (Fig. 6C). In this analysis we included 376 twin subjects (age: 29.2*±*3.4; 58.5% female) with data of both siblings available in both sessions. Session-specific heritability estimates are reported in Fig. S4.

For empirical data, heritability was 0.396*±*0.124 and 0.340*±*0.094 for SC and FC edges, and 0.349*±*0.110 and 0.358*±*0.077 for SC and FC node strengths, respectively. Simulated FC heritability was 0.275*±*0.098 at the level of edges and 0.256*±*0.088 at the level of nodes, which was significantly lower than empirical FC (edges: mean difference = -0.065, T = -25.75, p < 0.001; nodes: mean difference = -0.102, T = -8.73, p < 0.001). Simulated time-averaged state variables showed heritability estimates in comparable ranges 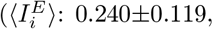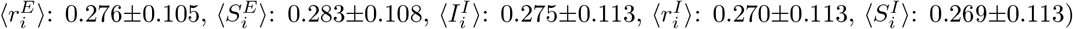. Given the influence of SC on simulated outcomes, we expected heritability patterns of simulated FC and state variables to align with those of SC. However, no such association was observed (Fig. S5).

Overall, while empirical features were more strongly heritable, simulated features also reflected genetic influences, indicating that BNMs can capture heritable aspects of brain function.

### 2.5 Scaling of GPU and CPU compute time with the number of simulations and network size

In the previous sections, we demonstrated the usage and applications of cuBNM, while comparing GPU and projected CPU compute times for those specific examples. Here, we systematically evaluated GPU and CPU performance for BNM simulations as a function of number of simulations and the network size. We compared run times on a single thread and all 72 cores (144 threads with hyperthreading) of a supercomputer CPU node using dual Intel Xeon IceLake Platinum 8360Y CPUs (2.4 MHz), with two Nvidia GPU models: the data-center-grade A100 (40 GB-SXM) and the consumer-grade GeForce RTX 4080 Super (16 GB).

We first investigated scaling of compute time with the number of simulations by running up to 2^15^ (32,768) simulations of rWW models with 100 nodes and identical parameters for 60 s (TR = 1 s, integration step = 0.1 ms). Except for the single-core CPU run, simulations were parallelized to the maximum hardware capacity. Running 32,768 simulations took 5.6 minutes on the A100 GPU, 21.8 minutes on the GeForce RTX 4080 Super, 82.4 minutes on the 72-core CPU node, and projected to take 3.8 days on a single CPU core. Relative to a single CPU core, the maximum speed-ups were 1,266.0x for the A100 GPU, 328.5x for the GeForce RTX 4080 Super, and 85.0x for the 72-core CPU node (Fig. 7A). Of note, the differences in scaling between the two GPU models and the multi-core CPU is due to their different parallelization capacity (number of physical or CUDA cores) as well as available memory, which influence the number of simulations that are calculated concurrently.

**Fig. 7.**
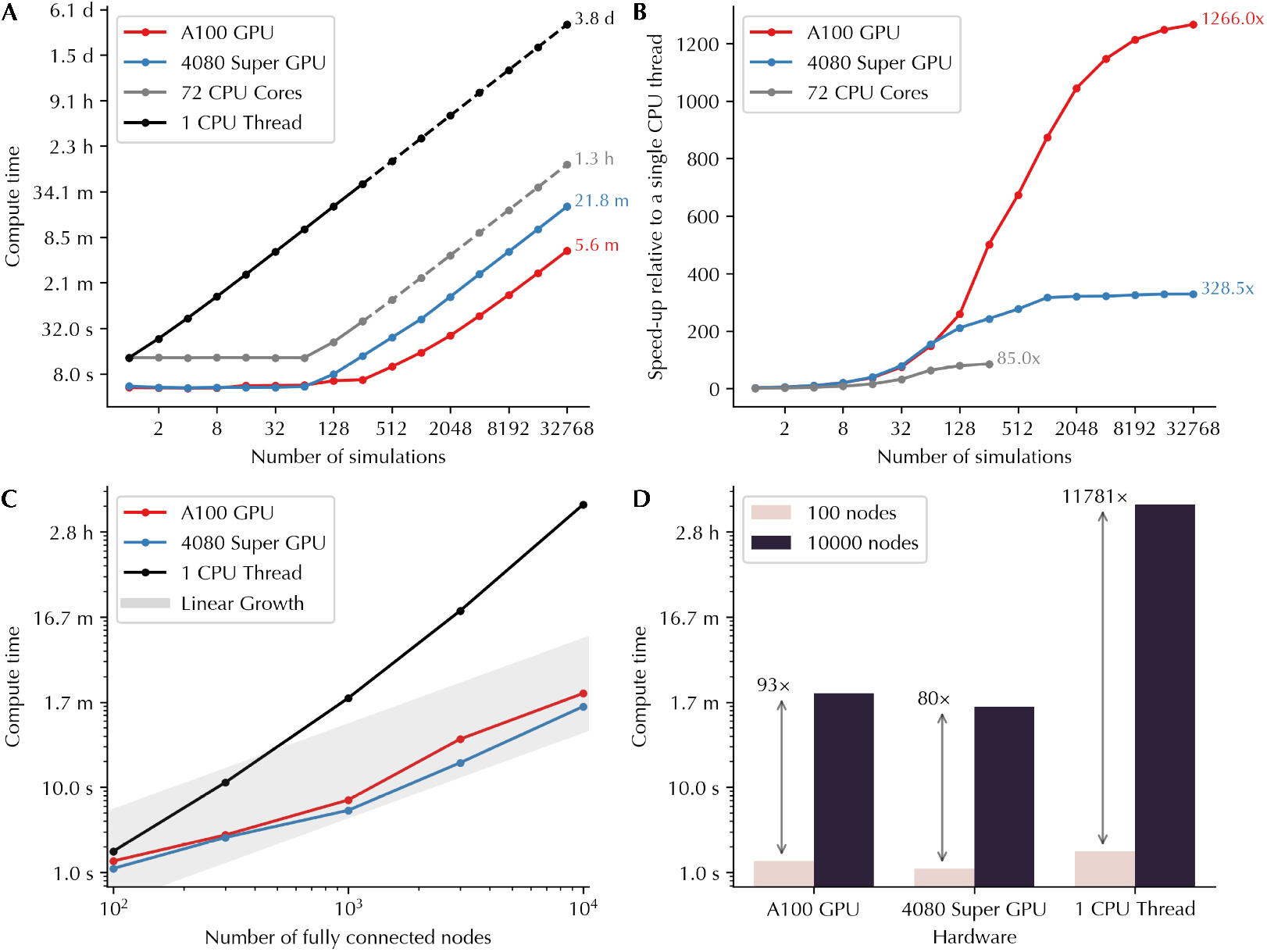
Scaling of GPU and CPU compute time with the number of simulations and network size. (**A**) Compute time as a function of the number of simulations (log-log scale). Solid lines show observed values and dotted lines indicate linear projections. (**B**) Speed-up relative to a single CPU thread as a function of the number of simulations (log scale on the x-axis). (**C**) Compute time as a function of network size (log-log scale). The grey band indicates the range of linear growth. GPU compute times increase linearly with the number of nodes, whereas CPU compute times increase supralinearly. (**D**) Comparison of compute times for simulations with 100 versus 10,000 nodes (log scale on the y-axis). Arrows indicate the fold-increase in compute time between the two network sizes.

Next, we assessed scaling with network size (number of nodes) by running a single rWW simulation with identical parameters for 60 s (TR = 1 s, integration step = 1 ms) using fully connected networks of 100, 300, 1,000, 3,000, and 10,000 nodes. We found that on both GPU models, compute time increased near-linearly with the number of nodes, but it increased supralinearly (approximately quadratically) on the CPU. For example, while running a 100-node simulation took 1.8 s on CPU and 1.4 s on the A100 GPU, increasing the network size by 100-fold to 10,000 nodes increased CPU run time to 20,849 s (11,781-fold increase) but A100 GPU run time to 126 s (93-fold increase) (Fig. 7B). Notably, this difference in scaling with network size is due to the implementation differences between CPU and GPU (Section 4.3.5). On the CPU, at each time step of each simulation, two nested loops over nodes exist: an outer loop that updates state of each node *i*, and an inner loop that computes the total input from all other nodes *j* to node *i*. This results in near-quadratic growth of compute time with the number of nodes. In contrast, on the GPU, the outer loop over nodes is parallelized across threads, leaving only the inner loop to be executed sequentially. Consequently, compute time on GPU grows near-linearly with the number of nodes. Notably, this near-linear scaling holds up to the maximum parallelization capacity of the device, beyond which simulations cannot be executed and an error is raised.

Overall, we observed that the GPU implementation of BNM simulations scaled efficiently with both the number of simulations and the number of nodes, offering massive performance gains compared to CPU implementations.

## 3 Discussion

In this study, we introduced cuBNM, a Python package with a C++ and CUDA backend that provides an efficient GPU-accelerated implementation of BNM simulations while also supporting CPUs. We demonstrated the usage of cuBNM and its bundled optimization approaches to fit both low-dimensional homogeneous and high-dimensional heterogeneous BNMs to group-level as well as individualized data. To illustrate the utility and biological relevance of individualized modeling, we performed model comparison, test-retest reliability, and heritability analyses in the HCP-YA twin dataset. We observed the node-based heterogeneous model outperforms both homogeneous and map-based heterogeneous models. Furthermore, within the node-based heterogeneous model, simulated features were found to be fairly reliable and heritable, highlighting biological relevance of individualized modeling.

Our scaling analyses showed that GPU parallelization leads to substantial performance gains relative to CPU implementations, with simulations running up to several hundred times faster. The massive speed-up of GPU-based implementation enables scalability of BNMs to larger samples, denser networks, and more complex models. Indeed, larger samples improve statistical power, can better capture population variance, and are critical for studies using machine learning. Furthermore, increasing network density and model complexity may offer additional value by improving biological realism of the model, which can in turn also improve model fit to the empirical data. Beyond computational speed, using GPUs additionally provides higher cost-efficiency, in terms of hardware costs, electricity consumption, and environmental impact. For example, in our individualized modeling analyses, we executed ∼ 28 million simulations taking ∼ 1,426 hours on the A100 GPU, corresponding to an estimated hardware rental cost of ∼ 1,556 € and electricity consumption of ∼ 1,027 kWh (carbon emission: ∼ 373 kg CO_2_). Equivalent multi-core CPU-based implementation was estimated to require ∼ 16,482 hours, corresponding to an estimated hardware rental cost of ∼ 48,504 €, and electricity consumption of ∼ 14,834 kWh (carbon emission: ∼ 5,385 kg CO_2_) leading to 11.5x slower compute time, 31.2x higher hardware costs, and 14.4x higher energy consumption and carbon emission (Supplementary Text A.2). The higher cost-efficiency of GPU-based implementation additionally makes the large-scale BNM studies more accessible, enabling smaller labs and institutes without supercomputing access to perform individualized modeling using a few consumer- or data-center-grade GPUs.

An additional advantage of efficient simulation frameworks is the facilitation of systematic evaluation of modeling choices. Previous studies have shown that BNM simulation outcomes can be influenced by the modeling and analytical choices^8,46,47^. Therefore, it is important for the BNM studies to demonstrate the robustness of their simulation-based findings to such variability^8^. The feasibility of repeated optimization under alternative configurations is substantially improved by GPU-based simulation, which is an important step toward enhancing robustness and replicability of simulation-based findings in the field.

In our individualized modeling analyses, we not only demonstrated the scalability and utility of cuBNM, but also highlighted several insights about BNMs and their simulated features. First, we found that heterogeneous models outperformed homogeneous models, largely due to higher *FC*_*corr*_. This finding is consistent with previous studies showing that introducing map-based heterogeneity improves model fit^8,9,12,36,42^. Extending these observations, we showed that node-based heterogeneous models further outperform map-based heterogeneous models. Of note, while map-based parameterization constrains the spatial distribution of parameters to the underlying maps and potentially introduces a bias, node-based parameterization allows greater flexibility in spatial variation across regions. Second, simulated features showed fair test-retest reliability (average ICC ≈ 0.40-50), while empirical FC was more reliable (average ICC ≈ 0.56-0.57). These estimates were in line with those reported in a prior study examining the reliability of simulated and empirical features, which also noted that in some modeling conditions, simulated features may show higher reliability than empirical ones^48^. We observed reliability was higher in sensorimotor regions, which were also characterized by stronger SC-FC coupling. This suggests that simulated measures are more stable in regions where FC is tightly constrained by SC, since the structural scaffold reduces variability and thereby increases reliability across repeated simulations which are fitted to the FC. Third, heritability analyses revealed *h*^2^ ≈ 0.25-0.30 for simulated features, compared to *h*^2^ ≈ 0.35-0.40 for empirical FC and SC. These estimates were comparable with prior heritability estimates reported for other structural^49,50^ and functional^51–53^ imaging traits, supporting the interpretation that simulated outcomes may capture biologically relevant individual differences that are partially grounded in genetic influences, and motivating future integration of BNMs with genome-wide association studies and imaging genetics. These example analyses illustrate the novel types of experiments that are made possible with the scalable individualized models. As an additional example, in our previous work^8^, we applied this GPU implementation to construct individualized heterogeneous BNMs for several hundred adolescents, and identified a replicable and robust maturational decrease of excitation-inhibition ratio in association cortical areas.

cuBNM is open-source, and we plan to continuously extend its functionality with community contributions. One important direction is extending the forward models to support simulating additional observation-level data such as magnetoencephalography/electroencephalography recordings^54,55^, as well as additional evaluation metrics, which can increase the applicability of cuBNM across different domains of brain imaging. In addition, while several widely used BNMs are currently implemented, facilitated by the modular design of the code, other models could be incorporated. For optimization, while grid search and various evolutionary optimizers from PyMoo are supported^38^, additional approaches such as Bayesian optimization^22^ and gradient-based methods^24,56^ would be valuable. In parallel to improving efficiency of numerical simulations, a complementary line of research focuses on estimating simulation outcomes without explicit numerical integration, using pre-trained machine learning models^57^. An intriguing opportunity for future research, is to leverage the efficient implementation of numerical simulations in cuBNM to generate large-scale training datasets across diverse modeling scenarios, which could facilitate the development and broader application of such learning-based approaches. Lastly, the core GPU implementation could be further extended to support GPU architectures beyond the currently supported models (Nvidia and AMD, the latter currently supported experimentally), in addition to enabling multi-GPU runs to further expand the parallelization capacity.

In conclusion, we showed that GPU-based implementation of BNM simulations provide substantial advantages in computational speed, cost-efficiency, and scalability, thereby enabling large-scale and individualized modeling. This GPU-based implementation offered through the cuBNM Python package, makes the BNM framework more accessible and feasible for a broader community of neuroscientists aiming to study diverse basic and clinical questions on mesoscale neural features inferred from macroscale dynamics. More broadly, as GPUs become increasingly widespread, there is a considerable promise for future research to accelerate a wider range of computationally intensive analyses in neuroscience and brain imaging by using GPUs^58,59^.

## 4 Materials and Methods

### 4.1 Participants

The Human Connectome Project - Young Adults dataset^30,31^ was used for all analyses presented in this study, as well as for the example data provided in cuBNM. A total of 1009 participants (age: 28.7*±*3.7; 54.1% female) with available T1-weighted (T1w), rs-fMRI, and DWI data were included. Depending on the specific analysis, we used either group-level or individual-level data from different subsets of this sample.

For group-level models, participants were divided into a training set (706 subjects; age: 28.7*±*3.7; 54.1% female) and a test set (303 subjects, age: 28.8*±*3.7; 54.1% female). The split was performed randomly while stratifying for sex and age group. For individualized models, we used the data of 430 twin participants (age: 29.2*±*3.3; 59.1% female), of whom 271 (62.9%) were monozygotic. In the main heritability analyses, we focused on 376 twin subjects (age: 29.2*±*3.4; 58.5% female) with complete functional data from both days 1 and 2 and both siblings. From the twins set, we randomly selected one member of each twin pair with available functional data from both sessions, to create a sample of 217 unrelated subjects (age: 29.3*±*3.4; 59.4% female), which were included in the test-retest reliability analysis. In addition, a random sample of 96 unrelated subjects (age: 29.2*±*3.2; 60.4% female) with available functional data from day 1 was selected from the twin subjects and was included in the model comparison analysis.

This research complies with the ethical regulations as set by The Independent Research Ethics Committee at the Medical Faculty of the Heinrich Heine University Düsseldorf (study number 2018-317).

### 4.2 Image acquisition and processing

The acquisition protocols of T1w, rs-fMRI and DWI scans are detailed in the HCP-YA documentation (https://www.humanconnectome.org/study/hcp-young-adult/document/1200-subjects-data-release)^30^. For the present analyses, it is of note that the rs-fMRI scans were acquired with a TR of 0.72 ms and a duration of 14:33 minutes per session.

#### 4.2.1 Functional image processing

We obtained the minimally-processed rs-fMRI data of the HCP-YA dataset^30,31^ in the *Cifti* space and parcellated it using the Schaefer-100 atlas^33^. We calculated FC as the correlation of BOLD signal time series between the cortical areas. In addition, dynamic FC matrices were calculated using sliding windows with a duration of 42 TRs and step size of 7 TRs, after discarding the edge windows. Next, the correlation of lower triangular parts of dynamic FC matrices was calculated between windows to generate the FCD matrix. Following Demirtas et al.^36^ and our previous work^8^, we excluded inter-hemispheric edges from the calculation of empirical and simulated FC and FCD matrices, to avoid detracting the optimization from fitting the network structure of FC within each hemisphere. Of note, excluding inter-hemispheric edges is not the default behavior in cuBNM, but the users have the option to include or exclude these connections.

Group-level FC matrices were obtained by Fisher Z-transforming individual FC matrices, averaging across sessions and subjects, and applying inverse Z-transformation. Group-level FCD distributions were derived using downsampled concatenation of individual distributions, first across sessions and then across subjects. Specifically, at each level, the lower triangular elements of individual FCD matrices were concatenated, sorted, and then subsampled at fixed intervals, leading to pooled FCD distributions that had a similar number of data points to the individual FCD distributions while preserving the underlying distribution. This was important to reduce memory and computational requirements of operations on group-level FCD distributions.

#### 4.2.2 Diffusion weighted image processing

DWI data was processed using an in-house pipeline^47^ (https://jugit.fz-juelich.de/inm7/public/vbc-mri-pipeline) that uses tools from *FreeSurfer*^60,61^, *FSL*^62^, *ANTs*^63^, and *MRtrix3* ^64^. Preprocessing included correction for eddy-current distortions and head motion, denoising, bias-field correction, and co-registration to the T1w image. Whole-brain tractography was performed in *MRtrix3* by first estimating fiber orientation distributions using spherical deconvolution, followed by probabilistic fiber tracking with 10 million streamlines using second-order integration over the fiber orientation distributions^65^. Next, the *tck2connectome* function of *MRtrix3* was used to reconstruct cortical node-wise SCs based on the Schaefer-100 atlas^33^. Group-averaged SCs were calculated for training and test sets by taking the median streamline counts for each edge across subjects. Finally, all SCs (individualized and group-averaged) were normalized by dividing each matrix by its mean × 100, resulting in a mean value of 0.01.

### 4.3 Brain network modeling

A BNM is created by connecting nodes of a network based on a connectivity matrix, and consists of (stochastic) differential equations that can model dynamic activity of brain nodes in interaction with each other and potentially external inputs. cuBNM is designed to construct BNMs that represent the target empirical data of a subject or a group. This process involves multiple steps, including: (i) definition of the network nodes and edges, (ii) mathematical description of the network dynamics, (iii) definition of free parameters and the homo-or heterogeneity of regional free parameters, (iv) generation of observation-level signal using a forward model, (v) numerical integration of the BNM and the forward model, (vi) computation of a cost function comprising of GOF and penalty terms, and (vii) model optimization using grid search or evolutionary optimizers. These steps are detailed in the subsections below.

#### 4.3.1 Defining the network nodes and edges

The brain network is defined as a weighted graph in which nodes represent brain areas, and edges represent the strength of connections between nodes. Nodes can be defined using existing anatomical, functional, or multimodal atlases at varying granularities. In this study, we used the Schaefer-100 cortical parcellation^33^. The edges are typically defined based on a group-level or individualized SC matrix derived from the DWI data, which quantifies the strength and pattern of white matter connections.

#### 4.3.2 Mathematical description of the network dynamics

Following, a mathematical description of the BNM needs to be specified, which governs the intrinsic (and stochastic) dynamics of each node as well as how they interact with each other. Various BNMs have been proposed in the literature, based on neural mass models or oscillators. We implemented several commonly used BNMs in cuBNM, including the excitatory-inhibitory rWW^35^, excitatory-only rWW^66^, Jansen-Rit^67^, Wilson Cowan^68,69^ and Kuramoto^70,71^ models (Table S1). Of note, a high-level YAML-based schema is used to define the models, including their variables, equations and metadata. This modular design enables researchers to more easily incorporate additional custom models in cuBNM.

In this study, we focused on the excitatory-inhibitory rWW model with FIC^35,36^. Here, each node consists of an excitatory (E) and an inhibitory (I) neuronal ensemble that interact with each other, modulated by a set of parameters: 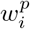 (local excitatory recurrence), 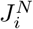 (NMDA receptor conductance), 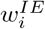 (inhibitory to excitatory connection weight) and *w*^*II*^ (local inhibitory recurrence). The nodes are connected to each other through their E neuronal ensembles according to the SC weights (*C*_*ij*_; connectivity strength of node *j* to node *i*) scaled by global coupling *G* as well as 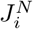. Additive Gaussian noise 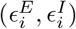 drives stochastic fluctuations and its magnitude is controlled by *σ*_*i*_. The node dynamics for a given node *i* is described by the following equations:

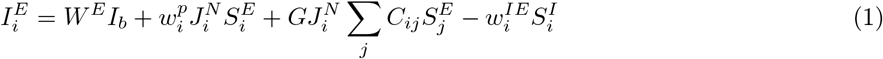

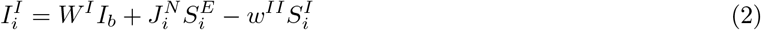

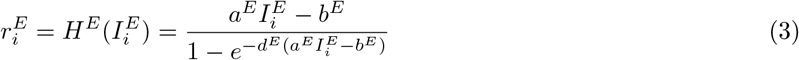

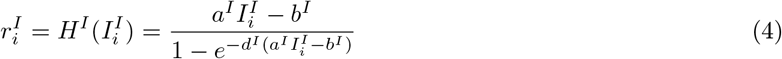

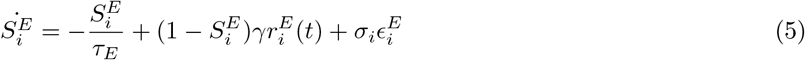

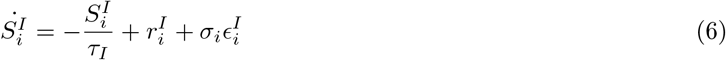

Here, 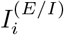 (input current [nA] to the E or I ensemble), 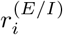 (firing rate [Hz] of E or I ensembles) and 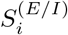 (synaptic gating variable of E or I ensembles) are the model *state variables* for each node *i*. Notably, 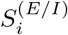 is restricted to the interval [0, 1]. The parameters used in the equations, along with their values, are summarized in Table S2.

##### Calculation of the global input to a node

The nodes in a BNM interact through the SC. At each simulation time point *t*, the total input to a *target* node *i* from all other *seed* nodes *j* (*j* ≠ *i*) is defined as the *global input*. For instance, in the rWW model, the global input corresponds to the term 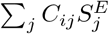 in Equation 1. Calculation of the global input requires specification of three components:

First, we define the state variable of the seed nodes that represents their outgoing signal, termed the *connectivity state variable x*. In the rWW model, this corresponds to 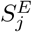.

Second, we specify how the connectivity state variables are combined. In conduction-based models such as rWW, this takes the form of a weighted summation 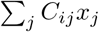, whereas in oscillator models such as Kuramoto, the global input is defined as the weighted sum of the phase differences between seed and target nodes, 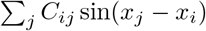.

Third, we determine whether there is conduction delay for the signal to reach from the seeds nodes to the target. In the absence of conduction delays (default in cuBNM and throughout the simulations reported in this study), the global input to node *i* at time *t* depends on the seed node activity at the immediately preceding time step *t* − *dt*. When conduction delays are included, node *i* receives input from node *j* at an earlier time point *t* − *τ*_*ij*_, where:

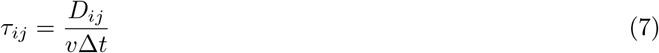

Here, *D*_*ij*_ is the white matter tract length between nodes *i* and *j* (mm), *v* is the conduction velocity (m/s), and Δ*t* is the integration step (ms). The delay *τ*_*ij*_ is expressed as the number of integration steps.

#### 4.3.3 Defining free parameters

cuBNM makes model parameters available as either potential free parameters or fixed constants. Notably, free parameters, except global coupling, are only *potentially* free, meaning that if not declared as a free parameter by the user, their value will be fixed to a default value. For the rWW model, potential free parameters include 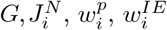, and σ_*i*_. We categorize the free parameters of each model into two broad groups: *global parameters*, which have a single global value for each simulation (e.g. *G*, and for simulations with conduction delay, *v*), and *regional parameters*, which in addition to having different values across simulations, can potentially have node-specific values (e.g. 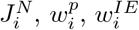, and 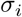) (Table S2). Notably, 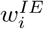 in rWW model with FIC is set based on the FIC algorithm to maintain 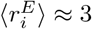 Hz (Supplementary Text A.3)^35,36^ and is therefore not tuned by the user or optimizer. In addition, in the present study *σ*_*i*_ was fixed to 0.01 nA and treated as a constant. Therefore candidate free parameters in this study were *G*, 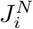, and 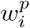.

Regional parameters within a simulation can be *homogeneous* (identical across nodes of a simulation) or *heterogeneous* (variable across nodes of a simulation). Heterogeneity can be specified via *map-based* or *node-based* approaches:

##### Map-based heterogeneity

Let *p* denote a heterogeneous regional parameter and *M* ∈ ℝ^*K × nodes*^ a matrix of *K* fixed node-wise maps. Regional values are defined by a parametric combination of maps with a bias *p*_*b*_ and coefficients 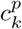:

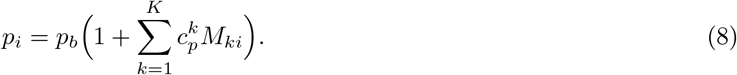

If any *p*_*i*_ falls outside predefined bounds, the global min and max are clipped to bounds and intermediate values are linearly redistributed.

In our demonstrations, we used two biological heterogeneity maps derived from the HCP-YA dataset: the T1w/T2w ratio map^31,72^ and the first principal gradient of FC (FC G1)^73^. These maps were obtained from the *neuromaps* package^74^, parcellated based on the Schaefer-100 atlas^33^, and min-max normalized to [0, 1]. cuBNM includes these and a few other example heterogeneity maps, but it also supports custom maps.

##### Node-based heterogeneous model

For a heterogeneous regional parameter *p*, the node-group parameters *p*_*g*_, *g* ∈ {1, …, *n*_*groups*_} become the tunable high-level model parameters which can be set independently from the other groups. Nodes that belong to the same group have the same parameter values. In our demonstrations, groups were defined based on the seven canonical resting-state networks^45^. Of note, custom groupings are supported, in addition to single-node groups (each node as its own group) or bilateral homologous groups (pairs of homologous nodes across hemispheres).

#### 4.3.4 Forward model

The neural activity of each node derived from the BNM, can subsequently be transformed to an observation-level signal using a forward model. Here, we used the hemodynamic model of Balloon-Windkessel to simulate node-wise BOLD signal. In each BNM, one of the state variables is selected to represent the neural activity of a given node. In the rWW model, 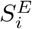 (*t*) is used for this purpose and serves as an input to the Balloon-Windkessel model, described by the following system of differential equations:

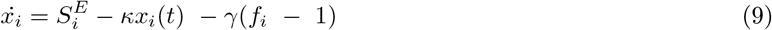

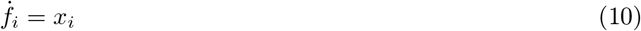

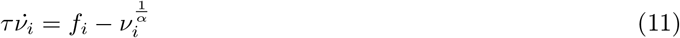

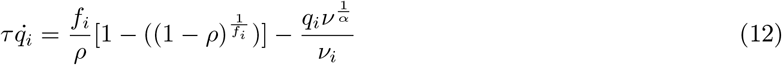

where *x*_*i*_ (vasodilatory signal), *f*_*i*_ (blood inflow), *ν*_*i*_ (blood volume) and *q*_*i*_ (deoxyhemoglobin content) are the model state variables in node *i*, and 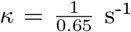 is the rate of signal decay, 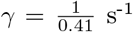 is the rate of flow-dependent elimination, *τ* = 0.98 s is the hemodynamic transmit time, α = 0.32 is the Grubb’s exponent, and *ρ* = 0.34 is the resting oxygen extraction fraction. Subsequently, the BOLD signal is calculated at every TR (0.72 s for the HCP-YA dataset) as:

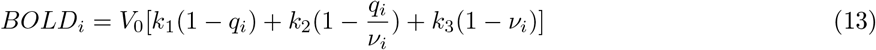

in which *V*_0_ = 2% is the resting blood volume fraction and *k*_1_ = 3.72, *k*_2_ = 0.527 and *k*_3_ = 0.53 are derived for 3T scans^36,75^.

#### 4.3.5 Numerical integration of the model equations

Having defined the network, the BNM, and the forward model, the next step is to run the simulation by numerically integrating the systems of differential equations for both the BNM and the Balloon-Windkessel model. The numerical integration of BNM is performed using the Euler-Maruyama approach, and the Balloon-Windkessel model is integrated using the Euler approach. The output of the simulation is the time series of BNM state variables as well as the BOLD signal derived from the Balloon-Windkessel model. In this study, all simulations had a duration of 900 s to match the duration of the functional scans in the HCP-YA dataset, except for the scaling analyses, which had a duration of 60 s.

In our implementation, the BNM and Balloon-Windkessel model are simulated simultaneously within a nested loop. By default, the Balloon-Windkessel model is simulated using an integration step of 1 ms, and BNMs are simulated using an integration step of 0.1 ms. Consequently, within each iteration of Balloon-Windkessel integration (outer loop), one or more BNM integration steps (inner loop, default = 10) are executed. The integration steps can be modified by the user. However, because of the nested design, the integration step of the Balloon-Windkessel model must be greater than or equal to that of the BNM.

##### Three levels of computations: simulations, nodes and time

Numerical integration requires evaluating the model equations at each simulation, node and time point. Across time, integration is necessarily serial, as current states depend on past states. However, across simulations and nodes, the calculations can be parallelized, subject to constraints. In the typical optimization workflow (Section 4.3.7), several simulations with different parameters are run. In cuBNM a *simulation group* (`SimGroup`) defines a set of simulations (with different parameters) intended for parallel execution to the extent of hardware capacity. For instance, the entire grid within the grid search, and each generation of an evolutionary optimizer, form a simulation group.

Within a simulation, calculations are performed for multiple nodes. However, except for an isolated network, the activity of each node in a BNM is influenced by the input from the other nodes. Therefore, the nodes are not completely independent, and their calculations cannot be fully parallelized for the whole duration of the simulation, but partial parallelization is possible. Specifically, calculations across nodes can be performed in parallel within each individual time step, after which nodes must be synchronized before continuing to the next integration step.

##### CPU implementation of simulations

On a single CPU thread, calculations for simulations, nodes, and time points are executed serially. With multiple threads, different simulations in a simulation group can run in parallel, using one thread dedicated to each simulation. Node-level parallelization is not applied on CPUs because of synchronization overhead, and given the number of network nodes often exceeds typical CPU thread counts.

##### GPU implementation of simulations

cuBNM includes two GPU implementation modes with different parallelization schemes. The *normal* mode is designed for running many parallel simulations of small-to-medium networks (< 500 nodes), and the *co-launch* mode supports running a few simulations of large networks (500 - several thousand nodes).

In the normal mode each simulation is assigned to a GPU “block”, and each node within a simulation is assigned to a GPU “thread” within the block of that simulation, and threads synchronize after each time step. For the large networks, given the hardware constraints on the number of threads within each block, this scheme is not suitable, and instead, the co-launch mode is used. In this mode, each simulation spans multiple blocks. The limitation of co-launch mode is that on most currently available GPU architectures, all blocks on the entire GPU must be synchronized together, which increases the synchronization overhead and reduces efficiency. Newer architectures with cluster-level synchronization can in principle improve efficiency for such cases. In addition, the maximum supported network size and number of simultaneous simulations are jointly constrained by the cooperative launch capacity of the device. This capacity limits the total number of blocks, which depends on both the number of nodes and simulations, and exceeding this limit results in a runtime error.

In addition, several technical considerations in the implementation of global input calculations should be noted. First, although intrinsic node dynamics are parallelized across threads, the computation of global input to each node requires iterating over all nodes within each thread. Consequently, computation time scales with network size, though substantially more efficiently than in CPU implementations, where a two-level nested loop over nodes is required. Second, the SC matrix is stored with sources as rows and targets as columns to ensure *coalesced memory access*, thereby reducing memory transfer burden and improving performance. Third, simulations with and without conduction delays differ in their implementation. Without conduction delays, only the connectivity state variable of the most recent time step needs to be stored. This limited memory demand allows storage within the *shared memory* specific to each GPU block, which is sufficient for small-to medium-sized networks. By contrast, simulations with conduction delays require storing a history of each node’s activity in a ring buffer, up to its maximum pairwise delay. This often exceeds the available shared memory capacity, necessitating the use of GPU global memory. As accessing global memory is slower than shared memory, this results in reduced computational efficiency for simulations with conduction delay.

##### Noise precalculation and shuffling

BNMs are typically stochastic and include noise terms, requiring Gaussian noise to be sampled for each node and time point. Sampling noise on-the-fly during simulation is computationally expensive. To address this, cuBNM precalculates the noise once on the CPU and reuses it across all simulations within a simulation group. Using a consistent noise input across simulations and subjects is advantageous for subject-specific fitting, as it prevents inter-individual differences in simulated outcomes from being driven by variation in the noise array. To reduce memory usage and CPU-to-GPU transfer overhead, a shorter noise segment (default: 30 s) is generated and repeated with shuffling throughout the simulation. To avoid spurious periodicity in simulated outcomes, these segments are independently shuffled across outer-loop iterations and nodes, based on the specified random seed.

#### 4.3.6 Model evaluation

Given a target empirical function data we define the cost function of a given simulation as:

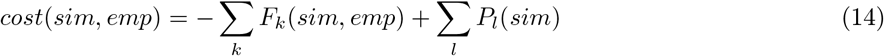

where *F*_*k*_ are GOF terms that quantify similarity of simulated and empirical data, and *P*_*l*_ are penalty terms independent of empirical data. Cost is computed after discarding the initial period (default: 30 s) of the simulation to ensure the BNM state has stabilized. The optimization goal is to minimize the cost function (i.e., maximize GOF and minimize penalty terms). In this paper, we defined cost function as:

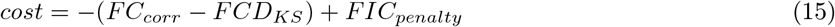

The details of each cost function term is described below:

##### Similarity of functional connectivity patterns

Simulated FC is calculated as the Pearson correlation between simulated BOLD time series of node pairs, excluding interhemispheric edges. It is computed efficiently on GPU (using custom CUDA kernels) or CPU (using *GNU Scientific Library*). Following, the correspondence of FC patterns is calculated as the Pearson correlation between the lower triangles of simulated and empirical FC matrices, between the empirical data and each simulation is evaluated by *FC*_*corr*_, with higher values indicating better fit. When available, GPU-based Pearson correlation from *cupy* package is used.

##### Similarity of functional connectivity dynamics

Simulated FCD is created by first calculating dynamic FC of sliding windows, followed by correlation of lower triangles of dynamic FC patterns between pairs of windows. It is computed efficiently on GPU (using custom CUDA kernels) or CPU (using *GNU Scientific Library*). Next, the dissimilarity of simulated and empirical FCD distributions is calculated as their Kolmogorov-Smirnov (KS) distance, *FCD*_*KS*_. When available, custom GPU kernels written in *numba-cuda* are used to calculated *FCD*_*KS*_. Because *FCD*_*KS*_ indicates dissimilarity, we use −*FCD*_*KS*_ as a GOF term.

##### Feedback inhibition control penalty

In the rWW model, despite the FIC algorithms are in place, 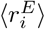 may diverge from the intended target of 3 Hz. Therefore, we define a *FIC*_*penalty*_ term to softly enforce the FIC target through optimization. We define:

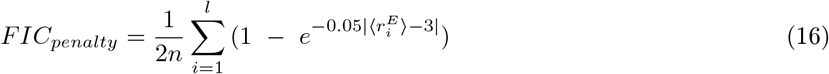

Here, the summation term is evaluated over *l* nodes with 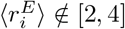 Hz, therefore node-wise penalty of nodes with 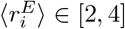 Hz is 0. The sum of node-wise penalties is then averaged over *n* (total number of nodes), and multiplied by 0.5, which is the 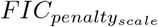 and determines the upper limit of *FIC*_*penalty*_. The presence and scaling of *FIC*_*penalty*_ can be modified by the user via 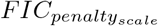 (`fic_penalty_scale`).

#### 4.3.7 Model optimization

The fitting of a BNM to target empirical data requires tuning or optimizing the model free parameters with the aim of minimizing cost function. In cuBNM two components are involved in model optimization:

1. BNM problem (`BNMProblem`) which characterizes the model type, network (nodes and their connectivity), free parameters (including their ranges, as well as the homo- or heterogeneity of regional parameters), simulation settings (e.g., duration, integration step, BOLD TR), and model evaluation metrics.
2. Optimizer (`Optimizer`) which is assigned to a BNM problem. It (iteratively) runs simulations with different combinations of free parameters, evaluates their cost function with respect to the provided empirical data, and ultimately identifies the optimal simulation with the lowest cost function. Two types of optimizers are available in cuBNM, as described below.

#### 4.3.8 Grid search

Grid search evaluates all combinations of evenly spaced parameter values within specified bounds and according to a specified resolution. This approach is practical for low-dimensional models. All grid simulations form a single simulation group which is executed in parallel up to hardware capacity.

#### 4.3.9 Evolutionary optimizers

Evolutionary optimizers iteratively sample the parameter space via generations of particles. Each particle represents a set of parameters, for which a simulation is run and a cost function is calculated. The cost functions of particles from each generation inform the sampling of the next, iteratively exploring the parameter space toward regions that minimize the cost. The particles in each generation form a simulation group which can be run in parallel. Notably, regardless of the hardware used for the execution of simulations and their evaluation against empirical data, the calculation of a new generation of sample solutions based on a given optimization algorithm is always done in CPU. cuBNM supports a large library of optimization algorithms that are implemented in the *PyMoo* Python package^38^.

##### Covariance Matrix Adaptation Evolution Strategy

In this study we used CMA-ES^39^, which performs well for fitting BNMs^8,9,42,76^. Each generation comprises Λ particles (here Λ = 128) exploring the parameter space collaboratively in an iterative process. After evaluating costs associated with the particles from each generation, the weighted mean of the best ⌊Λ*/*2⌋ particles defines the center of a multivariate normal distribution from which the next generation is sampled. The covariance is determined by a matrix which is updated to take the location of the current best points into account. Here, we set the iterations to repeat for up to 120 generations or until early termination, which occurs when the range of the best objective values over the last 10 + ⌈30*n/*Λ⌉ generations (where *n* is the number of tunable free parameters) as well as the current generation falls below *TolFun* = 0.005^39^. In individualized models, to reduce convergence to local optima, we performed each optimization run twice using different optimizer random seeds, and selected the better of the two optima.

##### Batch processing of evolutionary optimizers

In evolutionary optimization, the particles of a single run typically do not occupy the full parallelization capacity of the GPU, leading to its under-use and lower efficiency. In this case, to maximize GPU occupancy, cuBNM supports executing multiple evolutionary optimization runs (pertaining to different subjects and/or using different optimizer random seeds) concurrently on a single GPU.

### 4.4 Program structure

The cuBNM program is structured into two components (Fig. 8). First, the Python frontend provides the user interface via an API and a CLI, implements optimization and performs evaluation of simulations on CPU or GPU. Second, the core backend executes parallel simulations efficiently on the GPU or CPU. Given the simulation configuration and parameters received from the Python component, it calculates simulated BOLD, state variables, FC and FCD, and returns them to the Python component. The core consists of: (i) a C++ subcomponent that runs simulations and FC/FCD calculations on CPUs, while also acting as a bridge between Python and CUDA, and (ii) a CUDA subcomponent, that initializes arrays on GPUs, manages host-device memory transfers, and performs simulations as well as FC/FCD calculations on GPUs.

**Fig. 8.**
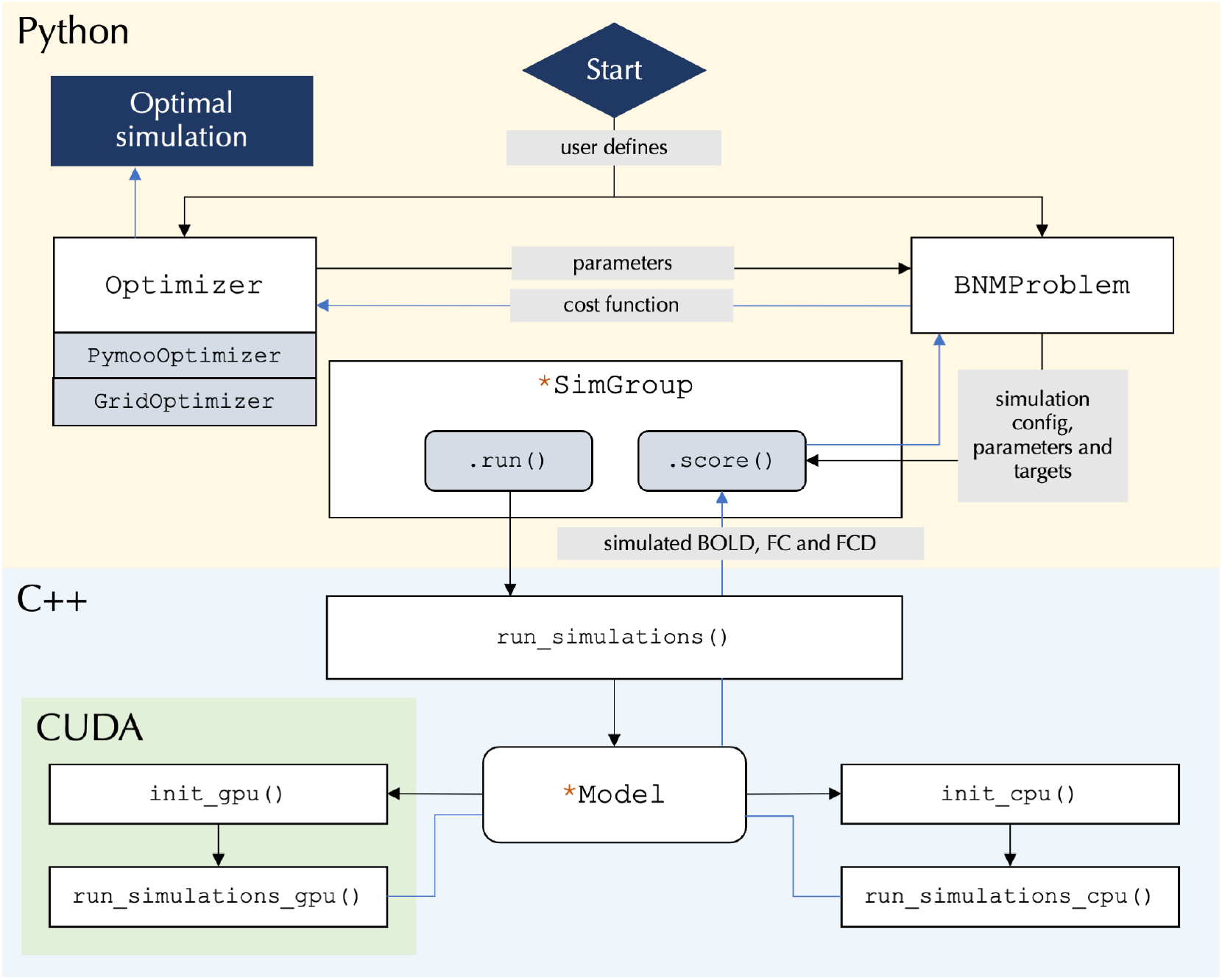
Program structure. Using the Python interface, the user defines a `BNMProblem` and assigns it to an `Optimizer` (of type `GridOptimizer` or `PymooOptimizer`). The `BNMProblem` is linked to a model-specific `SimGroup` (e.g., `rWWSimGroup`). `SimGroup.run()` calls the core `run_simulations`, which instantiates the corresponding C++ model (e.g., `rWWModel`), which runs the simulation on GPU/CPU. The core returns simulated BOLD and, when requested, FC and FCD, to the `SimGroup` instance, which scores them against empirical data via `SimGroup.score()`. The resulting cost function components are passed back to the `BNMProblem` and its `Optimizer`. In evolutionary optimization, these values guide the sampling of subsequent generations, with iterations continuing until the maximum generation count or an early termination criterion is reached. For grid search, a single exhaustive iteration is performed. In both cases, once the optimization completes, the optimal parameters are identified and returned to the user.

The Python API has four main modules:

- `cubnm.optimize` : Contains the `BNMProblem` class and the `Optimizer` family of classes. In typical workflows, these are the primary user-facing classes.
  - `BNMProblem` specifies a BNM optimization problem by defining the model type, network (nodes and their connectivity), free parameters (including their ranges, as well as the homo- or heterogeneity of regional parameters), simulation settings (e.g., duration, integration step, BOLD TR), and model evaluation metrics.
  - `Optimizer` includes `GridOptimizer` and `PymooOptimizer` family of classes (e.g., `CMAESOptimizer`). An optimizer is attached to a `BNMProblem` and (iteratively) tunes its parameters to minimize the cost function. This module also includes `batch_optimize()`, a function which enables performing multiple evolutionary optimizations in batch on a single GPU.
- `cubnm.sim` : Includes `SimGroup` and its model-specific subclasses (e.g., `rWWSimGroup`). `SimGroup` characterizes the network, simulation configuration, and parameters for a group of simulations that are intended to be run in parallel. In typical workflow, a `SimGroup` is created when a `BNMProblem` is instantiated, and users typically do not interact with it directly. `SimGroup.run()` calls `cubnm.core.run_simulations`, which is a backend function that runs simulations on CPU or GPU and returns simulated BOLD, state variables, FC/FCD and noise array. The outputs are passed to `SimGroup.score()` to compute the goodness-of-fit terms.
- `cubnm.datasets` : Contains loaders for the example data, such as structural connectomes, empirical BOLD, FC and FCD, as well as heterogeneity maps.
- `cubnm.utils` : Includes utility functions, notably GPU-accelerated Python functions for calculation of FC, FCD, *FC*_*corr*_, and *FCD*_*KS*_.

The full API is documented at https://cubnm.readthedocs.io/en/stable/autoapi/cubnm/index.html. In addition to the API, a CLI is installed with the package and accessible via the `cubnm` command (usage via `cubnm -h` ; details documented at https://cubnm.readthedocs.io/en/stable/cli.html). The CLI supports grid search (`cubnm grid`) and evolutionary optimization (`cubnm optimize`).

#### 4.4.1 Model definition in cuBNM

Model specification and implementation in cuBNM are facilitated through a code-generation framework based on YAML configuration files. This design enables developers and users to define and extend models in a structured manner. Briefly, each model is defined by a YAML file that specifies its metadata, the system of differential equations defining node dynamics (expressed as pseudo-code), details of global input computations, and the specification of model variables, including state variables, parameters, constants, and configuration options.

Based on these specifications, the framework automatically generates the corresponding Python, C++, and CUDA code required for simulation and integration within the toolbox. This approach abstracts away low-level implementation details and facilitates addition of new models. In addition, advanced use cases are supported, allowing contributors to provide custom C++/CUDA code snippets to override or extend specific components of the simulation pipeline. Of note,the code-generation process occurs at compile time. Therefore, newly defined models require rebuilding the package from source, and cannot be introduced dynamically at runtime. A detailed guide for adding new models is provided at https://cubnm.readthedocs.io/en/stable/contribution.html.

### 4.5 Statistical analyses

#### Test-retest reliability

We evaluated test-retest reliability of each feature using functional scan sessions acquired on day 1 and day 2 in a cohort of 217 unrelated HCP participants. Reliability was quantified using the intraclass correlation coefficient^77^ ICC(2,k), implemented via the *pingouin* package^78^. For node-wise and edge-wise features, the ICC was calculated independently for each node or edge. The ICC values were interpreted as follows: <0.40, poor; between 0.40 and 0.60, fair; between 0.60 and 0.75 good; above 0.75, excellent^79^.

#### Heritability

We estimated heritability based on the data from the twin subjects of the HCP dataset and in line with our previous work^50,52,53,80,81^. Heritability estimates and their standard errors were obtained with Sequential Oligogenic Linkage Analysis Routines (SOLAR, version 9.0.0), which applies maximum likelihood variance decomposition to partition phenotypic covariance into familial and environmental components, by modeling the covariance among family members as a function of genetic proximity^82^. Narrow-sense heritability (*h*^2^) was defined as the proportion of total phenotypic variance 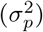 explained by additive genetic variance 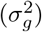, that is 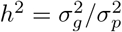. Inverse normal transformation was applied to features, and each feature was modeled using the A + E model, using age, sex, age^2^, and age*sex as the confounds^83^. We used the GPU version of fast permutation heritability inference approach for heritability calculations^49^. For features derived from simulated and empirical functional data with two available sessions, heritability was estimated on values averaged across sessions. Session-specific heritability values were also computed and are presented in Fig. S4.

#### Spin test

We assessed the spatial association between a source map *X* and a target map *Y* using Pearson correlation. To account for spatial autocorrelation, we generated non-parametric null distributions of correlation coefficients by permuting the target map using surface-based spin rotations. Spin permutations were implemented at the parcel level using the *neuromaps* package^74^, following the method of Alexander-Bloch et al.^84^. For each test, 1000 permutations were performed.

### 4.6 Scaling analyses

We evaluated compute time as a function of both the number of simulations and the network size (number of nodes). Analyses were performed on four hardware configurations: (i) a single thread or (ii) all 72 cores (144 threads with hyperthreading) of a supercomputer CPU node using two inter-connected Intel Xeon IceLake Platinum 8360Y CPU (2.4 GHz), (iii) a single Nvidia A100 GPU (40 GB-SXM), (iv) a single GeForce RTX 4080 Super GPU (16 GB). Each scaling experiment was run twice, and the average of the two compute times was reported.

To assess the scaling of compute time with the number of simulations, we ran simulations of rWW model with FIC using 100 nodes connected based on the group-averaged SC of HCP training subjects. Each simulation was 60 s long, with TR = 1 s and integration step = 0.1 ms. Identical parameters were used across all simulations. Except for single-threaded CPU runs, simulations were parallelized to the full extent of hardware capacity. The number of simulations ranged from 1 up to 2^15^ (32,768) on GPUs, and 1 up to 2^8^ (256) on CPUs. For multi-core and single-thread CPU, the projected compute time for larger number of simulations was estimated by linear extrapolation from the last measured data point. Given compute time Δ*t* on GPU or multi-core CPU, speed-up values were calculated relative to the single-thread CPU compute time Δ*t*_*single*_ as *N*_*simulations*_*/*(Δ*t*/Δ*t*_*single*_).

To assess scaling with network size, we executed a single rWW simulation using fully connected networks of 100, 300, 1,000, 3,000, and 10,000 nodes. Each simulation was 60 s long, with TR = 1 s and integration step = 1.0 ms. On GPUs, simulations were performed in co-launch mode to enable simulation of denser networks, which is not feasible in normal mode.

### 4.7 Data availability

The HCP-YA dataset is openly accessible to registered researchers at https://www.humanconnectome.org/study/hcp-young-adult, and is also available as a DataLad dataset at https://github.com/datalad-datasets/human-connectome-project-openaccess. Heterogeneity maps were obtained from the *neuromaps* Python package.

### 4.8 Code availability

The source code for cuBNM is openly available on GitHub (https://github.com/amnsbr/cubnm) under the BSD 3-Clause License. This code is also archived on Zenodo (https://doi.org/10.5281/zenodo.12097797), with the DOI corresponding to the most recent release. cuBNM can be installed as a Python package through PyPi (https://pypi.org/project/cubnm) and is additionally available as a Docker container (https://hub.docker.com/r/amnsbr/cubnm). Comprehensive documentation and tutorials, including details of the API, are provided at https://cubnm.readthedocs.io/en/stable.

The code for the analyses reported in this manuscript is available on GitHub (https://github.com/amnsbr/cubnm_paper) and is deposited on Zenodo (https://doi.org/10.5281/zenodo.17630927). All analyses were conducted using development versions of cuBNM between stable releases v0.0.7 and v0.1.0. Heritability analyses were performed using SOLAR (version 9.0.0 dynamic; www.nitrc.org/projects/se_linux/). DWI preprocessing was performed using an in-house pipeline available at https://jugit.fz-juelich.de/inm7/public/vbc-mri-pipeline.

## 5 Acknowledgments

AS, BW and SLV were funded by the Max Planck Society. AS and SLV were additionally funded by Helmholtz Association’s Initiative and Networking Fund under the Helmholtz International Lab grant agreement InterLabs0015, and the Canada First Research Excellence Fund (CFREF Competition 2, 2015-2016) awarded to the Healthy Brains, Healthy Lives initiative at McGill University, through the Helmholtz International BigBrain Analytics and Learning Laboratory (HIBALL). SBE was supported by the Deutsche Forschungsgemeinschaft (DFG, EI 816/21-1), the National Institute of Mental Health (R01-MH074457), and the European Union’s Horizon 2020 Research and Innovation Programme under Grant Agreement No. 945539 (HBP SGA3). BCB acknowledges research support from the National Science and Engineering Research Council of Canada (NSERC RGPIN-2025-05932), CIHR (FDN-154298, PJT-174995, PJT-191853, PJT-203761), SickKids Foundation (NI17-, HIBALL, Healthy Brains and Healthy Lives (HBHL), Brain Canada Foundation, FRQS, the Tier-2 Canada Research Chairs Program, and the Centre for Excellence in Epilepsy at the Neuro (CEEN). The authors acknowledge computing time on the supercomputer JURECA^85^ at Forschungszentrum Jülich under grant no. “eidev”. They additionally acknowledge computing time on the supercomputer Raven at Max Planck Computing and Data Facility, as well as high throughput computing cluster Juseless at Forschungszentrum Jülich. Language editing of this manuscript was assisted by large language models.

## Footnotes

a In the literature, also known as “biophysical network modeling” or “whole-brain dynamical modeling”. Notably, dynamical causal modeling, a specific type of biophysical network modeling that focuses on the estimation of effective connectivity using a Bayesian framework, is not the subject of this study.

