## Supplementary for "cuBNM: GPU-Accelerated Brain Network Modeling"

Saberi et al.

### A Supplementary Text

#### A.1 Identity of simulations calculated on CPU and GPU

Here, we assessed the identity of simulated outputs calculated on CPU compared to GPU. We performed a 3D grid search of the homogeneous reduced Wong-Wang (rWW) model, sampling each parameter $G$, $w^p$, and $J^N$ at 10 evenly spaced values. Each simulation was run for 60 s (TR = 1 s). The mean absolute difference between CPU and GPU simulations was on the order of $10^{-16}$ to $10^{-14}$ across blood-oxygen-level-dependent (BOLD) signals, FC, FCD, and state variables, which is negligible and within the limits of double-precision floating-point arithmetic. The grids of negative cost values, as well as an example BOLD time series calculated on CPU and GPU, are shown in Fig. S6. These results demonstrate a near-perfect identity between CPU and GPU simulations of the rWW model across a wide parameter range.

Asserting the identity of CPU and GPU outputs for an example parameter set constitutes one of the unit tests performed in the program for every included model. Of note, while CPU and GPU outputs are identical across all included models, exceptions can occur in the Kuramoto model, where this identity does not strictly hold. The Kuramoto model shows parameter-dependent divergence between CPU and GPU implementations (particularly with high coupling, delays, or noise), likely due to its sensitivity to small numerical differences that amplify through model dynamics. This reflects discrepancies regarding the exact implementation and potential rearrangement of mathematical operations rather than computational errors, as independent implementations (e.g., a Python CPU-based implementation) also produce distinct trajectories under these conditions. For more discussion on this topic, we refer to the relevant issue in cuBNM GitHub repository (`https://github.com/amnsbr/cubnm/issues/24`).

#### A.2 Estimation of computational time, hardware cost, energy expenditure and carbon footprint of simulations run on GPUs versus CPUs

In our individualized modeling analyses, we performed a total of 28,158,401 simulations, taking 1,426.0 hours on Nvidia A100 GPUs. We estimated the compute time on a multi-core CPU node (two Intel Xeon IceLake Platinum 8360Y CPUs; 72 cores; 2.4 GHz) by running 128 parallel simulations, corresponding to one generation of the optimization runs. The compute time for running and scoring each simulation on this CPU node was 2.15 s, therefore projecting the full set of simulations to take a total of 16,842.0 hours, which is 11.8 times the compute time observed on GPUs.

For hardware cost estimation, as of September 2025, we obtained an hourly rental price of 1.29 $ (1.09 €) for the Nvidia A100 GPU (40GB-SXM) from a commercial provider. No rental price was found for the specific CPU model tested. Instead, we used a generic dedicated Intel CPU rental price of 0.04 € per core per hour, obtained from a separate provider, which corresponds to 2.88 € per hour for a 72-core node. To estimate electricity consumption, we multiplied the thermal design power (TDP) specified by the manufacturer (400 W for the GPU; 250 W per CPU for the dual-CPU node, i.e., 500 W per node) by the power usage effectiveness set to 1.8, according to an average reported in 2020[1]. The resulting total electricity consumption was then used to estimate $CO_2$ emissions by multiplying with the average grid carbon intensity in Germany in 2024 (363 g $CO_2$/kWh)[2].

#### A.3 Feedback inhibition control in the reduced Wong-Wang model

The feedback inhibition control (FIC) algorithm in the rWW model determines the regional values of $w_i^{IE}$ given the structural connectome (SC) and the other model parameters, aiming to maintain an average excitatory firing rate close to 3 Hz, which is proposed to be the biological range in a state of excitation-inhibition balance[3]. Both analytical[4] and numerical[3] implementations of the FIC algorithm are available in cuBNM. By default, cuBNM only uses the analytical implementation. However, it can be configured to additionally perform numerical adjustments using a specific number of numerical FIC trials, with the initial $w_i^{IE}$ values being set to the analytical estimates.

**Analytical implementation.** This implementation proposed by Demirtaş et al.[4] analytically solves for $w_i^{IE}$ to satisfy the self-consistency of the model equations at the steady state with $\langle r^E \rangle \approx 3$ Hz, corresponding to $\langle S^E \rangle \approx 0.164757$ and $\langle I^E \rangle \approx 0.37738$ nA, following:

$$w_i^{IE} = \frac{W^E I_b + w_i^p J_i^N \langle S^E \rangle + G J_i^N \sum_j C_{ij} \langle S^E \rangle - \langle I^E \rangle}{\langle S_i^I \rangle} \tag{17}$$

Here, the steady-state $\langle S_i^I \rangle = H^I(\langle I_i^I \rangle)\tau^I$ is estimated by solving for $\langle I_i^I \rangle$ in:

$$W^I I_b + J_i^N \langle S^E \rangle - H^I(\langle I_i^I \rangle)\tau^I - \langle I_i^I \rangle = 0 \tag{18}$$

**Numerical implementation.** The numerical implementation[3] iteratively refines $w_i^{IE}$ by directly integrating the model equations. Starting from the analytical estimates, the system is simulated for 10 s, after which the mean excitatory input current $\langle I_i^E \rangle$ for each node is computed over the interval from 1 s to 10 s. If the difference $\langle I_i^E \rangle - \frac{b_E}{a_E}$ deviates from the target value of $-0.026$ nA by more than 0.005 nA, $w_i^{IE}$ is increased/decreased when input current is higher/lower than the target. The simulation is then repeated with the updated parameters. This iterative adjustment continues for the user-defined number of trials or until all nodes reach the FIC target. Of note, unlike the original implementation where all $w_i^{IE}$ values were initialized to 1, in cuBNM the initial $w_i^{IE}$ is set to the analytical estimates.

### B Supplementary Tables

Table S1: List of implemented models

| Model | Equations | Free parameters[a] | Connectivity state variable | BOLD state variable |
|---|---|---|---|---|
| Reduced Wong-Wang[3] ( `rWW` ) | $I_i^E = W^E I_b + w_i^p J_i^N S_i^E + G J_i^N \sum_j C_{ij} S_j^E - w_i^{IE} S_i^I$<br>$I_i^I = W^I I_b + J_i^N S_i^E - w^{II} S_i^I$<br>$r_i^E = H^E(I_i^E) = \frac{a^E I_i^E - b^E}{1 - e^{-d^E(a^E I_i^E - b^E)}}$<br>$r_i^I = H^I(I_i^I) = \frac{a^I I_i^I - b^I}{1 - e^{-d^I(a^I I_i^I - b^I)}}$<br>$\dot{S_i^E} = -\frac{S_i^E}{\tau_E} + (1 - S_i^E)\gamma r_i^E(t) + \sigma_i \epsilon_i^E$<br>$\dot{S_i^I} = -\frac{S_i^I}{\tau_I} + r_i^I + \sigma_i \epsilon_i^I$ | $G$, $w_i^p$, $J_i^N$, $w_i^{IE}$, $\sigma_i$ | $S_i^E$ | $S_i^E$ |
| Reduced Wong-Wang (Excitatory)[5] ( `rWWExc` ) | $x_i = w_i J^N S_i + G J^N \sum_j C_{ij} S_j + I_{0_i}$<br>$r_i = \frac{a x_i - b}{1 - e^{-d(a x_i - b)}}$<br>$\dot{S}_i = -\frac{S_i}{\tau_s} + (1 - S_i)\gamma r_i + \sigma_i \epsilon_i$ | $G$, $w_i$, $I_{0_i}$, $\sigma_i$ | $x_i$ | $x_i$ |
| Jansen-Rit[6] ( `JR` ) | $\dot{y_{0_i}} = y_{3_i}$<br>$\dot{y_{1_i}} = y_{4_i}$<br>$\dot{y_{2_i}} = y_{5_i}$<br>$\dot{y_{3_i}} = Aa[\frac{2\nu_{max}}{1+e^{r(v_0 - (y_{2_i} - y_{1_i}))}}] - 2a y_{3_i} - a^2 y_{0_i} + \sigma_i \epsilon_i$<br>$\dot{y_{4_i}} = Aa[\mu + a_{2_i} J_i \frac{2\nu_{max}}{1+e^{r(v_0 - (a_{1_i} J y_{1_i}))}} + \sum_j C_{ij} x_j$<br>$\dot{y_{5_i}} = Bb(a_{4_i} J_i [\frac{2\nu_{max}}{1+e^{r(v_0 - (a_{3_i} J y_{0_i}))}}]) - 2b y_{5_i} - b^2 y_{2_i}$<br>$z_i = y_{1_i} - y_{2_i}$<br>$x_i = c_{min} + \frac{c_{max} - c_{min}}{1+e^{r(c_{mid} - z_i)}}$ | $G$, $J_i$, $a_{1_i}$, $a_{2_i}$, $a_{3_i}$, $a_{4_i}$, $\sigma_i$ | $x_i$ | $z_i$ |
| Wilson-Cowan[7,8] ( `WC` ) | $\tau^E \dot{E}_i = -E + (1-E) S^E [c^{EE} E_i - c^{IE} I_i + G \sum_j C_{ij} E_j + P_i^E] + \sigma_i^E \epsilon_i^E$<br>$\tau^I \dot{I}_i = -I + (1-I) S^I [c^{EI} E_i - c^{II} I_i + P_i^I] + \sigma_i^I \epsilon_i^I$, where:<br>$S^E(x) = \frac{1}{1+e^{-a^E(x-\mu^E)}}$<br>$S^I(x) = \frac{1}{1+e^{-a^I(x-\mu^I)}}$ | $G$, $c_i^{EE}$, $c_i^{EI}$, $c_i^{IE}$, $c_i^{II}$, $P_i^E$, $P_i^I$, $\sigma_i^E$, $\sigma_i^I$ | $E_i$ | $E_i$ |
| Kuramoto[9,10] ( `Kuramoto` ) | $\dot{\theta}_i = \omega_i + G \sum_j C_{ij} \sin(\theta_j - \theta_i) + \sigma_i \epsilon_i$ | $G$, $\omega_i$, $\sigma_i$, $\theta_{init_i}$ | $\theta_i$ | $\theta_i$ |

[a] In all models, when conduction delay is introduced, conduction velocity $v$ is an additional global free parameter.

Table S2: Reduced Wong-Wang model terms

| Term | Description | Value |
|---|---|---|
| $I_i^E$ | Input current to $E$ pool | State variable |
| $I_i^I$ | Input current to $I$ pool | State variable |
| $r_i^E$ | Firing rate of $E$ pool | State variable |
| $r_i^I$ | Firing rate of $I$ pool | State variable |
| $S_i^E$ | Synaptic gating variable of $E$ pool | State variable |
| $S_i^I$ | Synaptic gating variable of $I$ pool | State variable |
| $G$ | Global coupling | Global free parameter |
| $w_i^p$ | Local excitatory recurrence | Regional free parameter |
| $J_i^N$ | NMDA receptor conductance (nA) | Regional free parameter |
| $\sigma_i$ | Gaussian noise amplitude (nA) | Regional free parameter[a] |
| $I_b$ | Baseline input current (nA) | 0.382 |
| $W^E$ | Weight of baseline input current to $E$ pool | 1.0 |
| $W^I$ | Weight of baseline input current to $I$ pool | 0.7 |
| $w^{II}$ | Weight of recurrent inhibitory connections | 1.0 |
| $H^E$ | Input-output function of $E$ pool | - |
| $a^E$ | Slope of $H^E$ (nC) | $310^{-1}$ |
| $b^E$ | Threshold above which $E$ firing rate increases linearly with the input current (nA) | 0.403 |
| $d^E$ | Shape of $H^E$ curvature around $b^E$ | 0.16 |
| $H^I$ | Input-output function of $I$ pool | - |
| $a^I$ | Slope of $H^I$ (nC) | $615^{-1}$ |
| $b^I$ | Threshold above which $I$ firing rate increases linearly with the input current (nA) | 0.288 |
| $d^I$ | Shape of $H^I$ curvature around $b^I$ | 0.087 |
| $\tau_E$ | NMDA receptor decay time constant (s) | 0.1 |
| $\gamma$ | NMDA kinetics rate constant | 0.000641 |
| $\tau_I$ | GABA receptor decay time constant (s) | 0.01 |
| $\epsilon_i^E$ | Gaussian noise input to $E$ pool | Sampled from $\mathcal{N}(0,1)$ |
| $\epsilon_i^I$ | Gaussian noise input to $I$ pool | Sampled from $\mathcal{N}(0,1)$ |

[a] In this manuscript, $\sigma_i$ is fixed to 0.01 nA in all simulations and nodes. However, in cuBNM it can be set as a free parameter.

Table S3: Summary of optimal model fits across optimization approaches and model types in group-level models.

| Model | Optimization | Training | | | | Test | | | |
|---|---|---|---|---|---|---|---|---|---|
| | | Cost | $FC_{corr}$ | $FCD_{KS}$ | $FIC_{penalty}$ | Cost | $FC_{corr}$ | $FCD_{KS}$ | $FIC_{penalty}$ |
| Homogeneous | Grid search | -0.119 | 0.298 | 0.152 | 0.028 | -0.118 | 0.297 | 0.152 | 0.028 |
| Homogeneous | CMA-ES | -0.125 | 0.299 | 0.140 | 0.034 | -0.124 | 0.298 | 0.140 | 0.034 |
| Map-based heterogeneous | CMA-ES | -0.267 | 0.511 | 0.198 | 0.044 | -0.258 | 0.496 | 0.194 | 0.044 |
| Node-based heterogeneous | CMA-ES | -0.338 | 0.521 | 0.170 | 0.013 | -0.327 | 0.508 | 0.168 | 0.014 |

$FC_{corr}$: functional connectivity correlation; $FCD_{KS}$: functional connectivity dynamics Kolmogorov–Smirnov distance; $FIC_{penalty}$: functional inhibition control penalty

### C Supplementary Figures

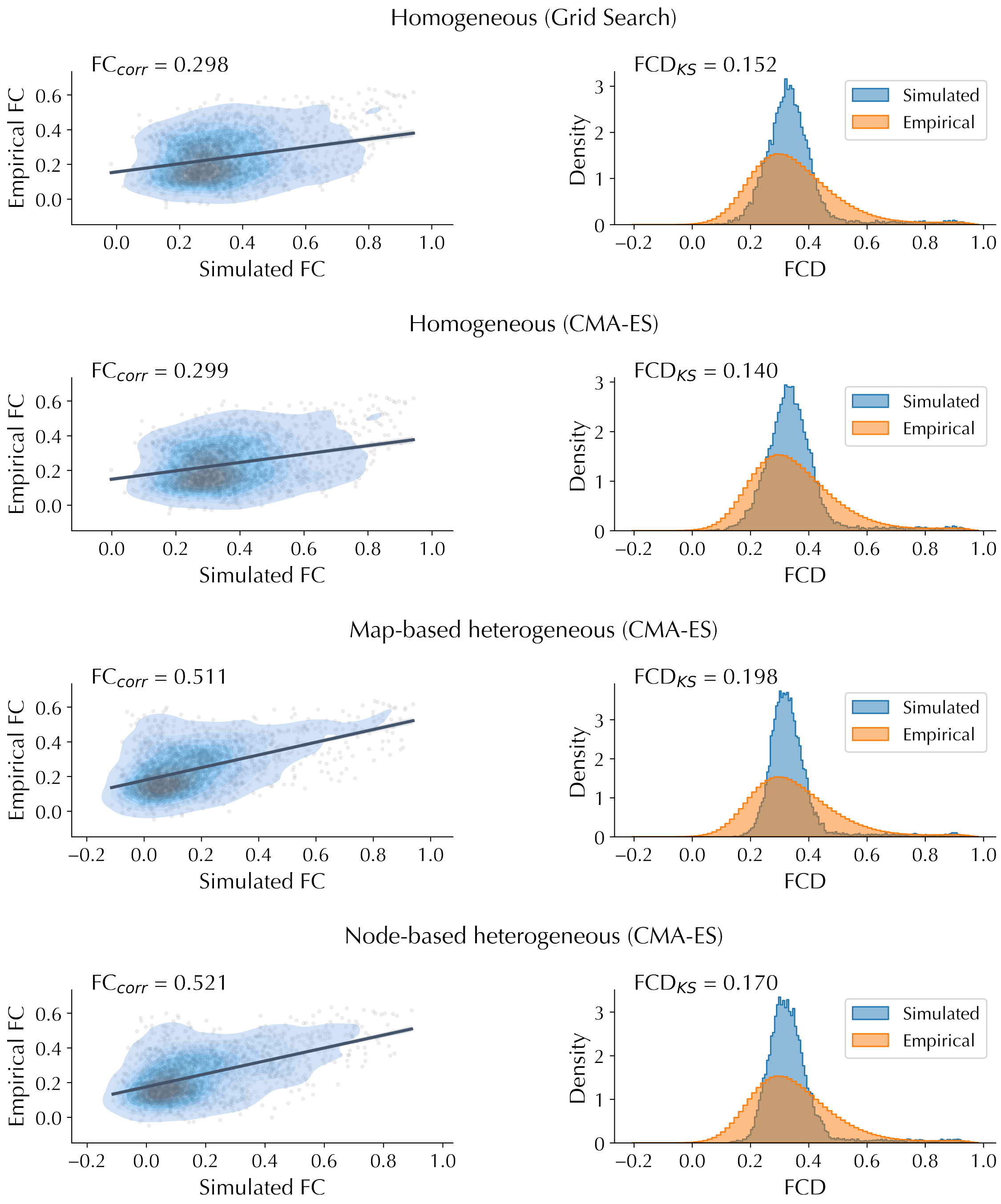


**Fig. S1. Fit of optimal simulations to empirical data in group-level models.** The correspondence between simulated and empirical functional connectivity (FC) (*left*) and the distributions of simulated and empirical functional connectivity dynamics (FCD) for each model and optimization approach is shown.

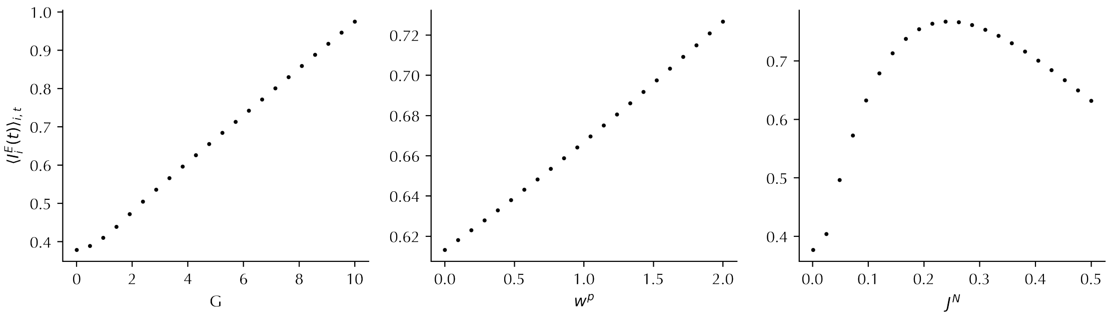


**Fig. S2. Variation of average excitatory input current as a function of model parameters.** Each plot shows how the average input current to excitatory neurons $\langle I_i^E(t)\rangle_{i,t}$ varies as a function of the indicated model parameter. For each parameter, values were averaged across all $22^2 = 484$ simulations with that parameter value, collapsing across the other two parameters.

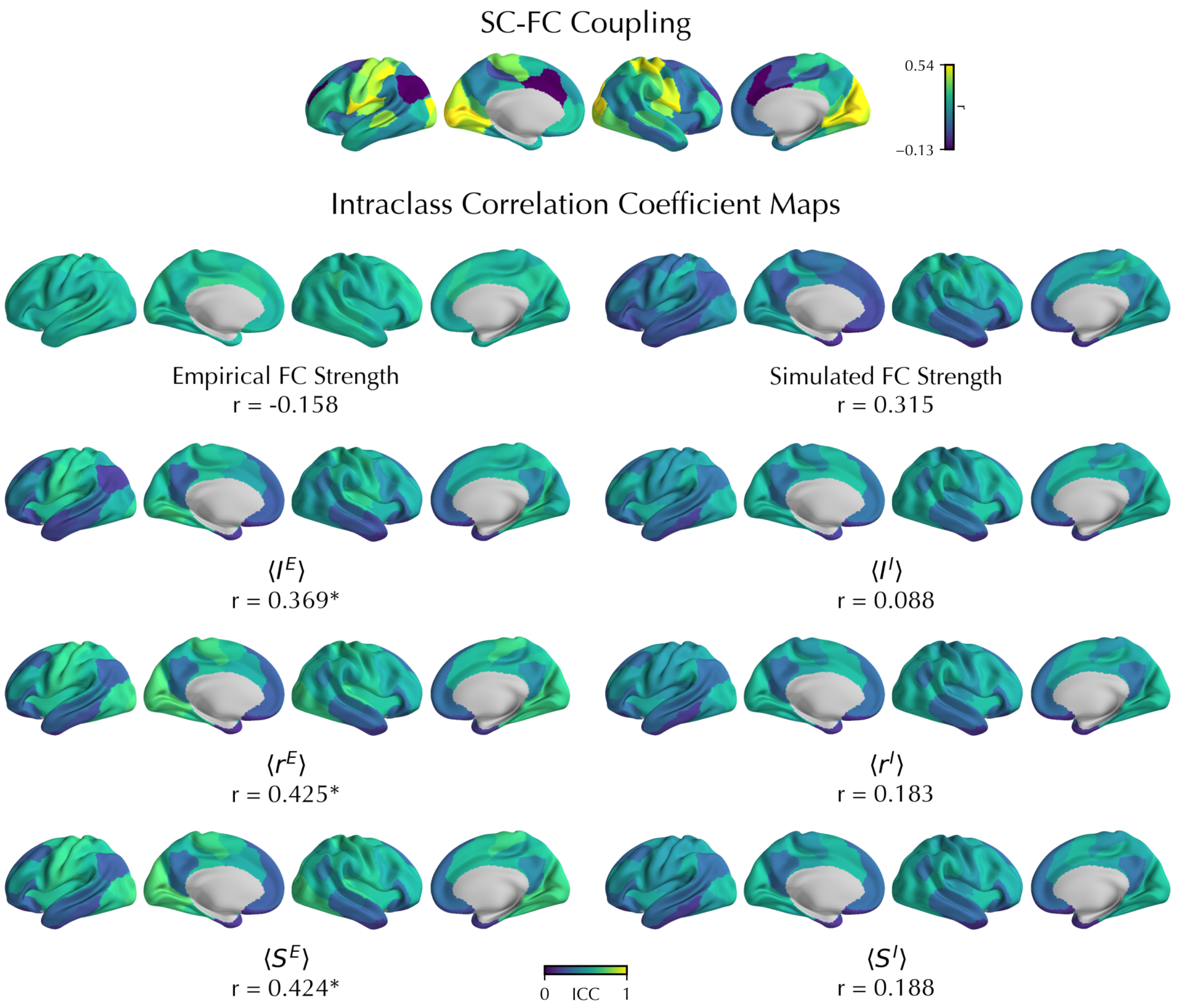


**Fig. S3. Spatial pattern of test-retest reliability in relation to SC-FC coupling.** *Top*: Regional values of group-averaged coupling between the structural connectome (SC) and functional connectome (FC), calculated as the row-wise Pearson correlation coefficient and averaged across sessions and subjects. *Bottom*: Regional intraclass correlation coefficients (ICCs) for empirical and simulated measures derived from day 1 and day 2 scans. Reported statistics below each map indicate the Pearson correlation between the ICC map and the SC-FC coupling map. Asterisks denote $p_{spin,FDR} < 0.05$.

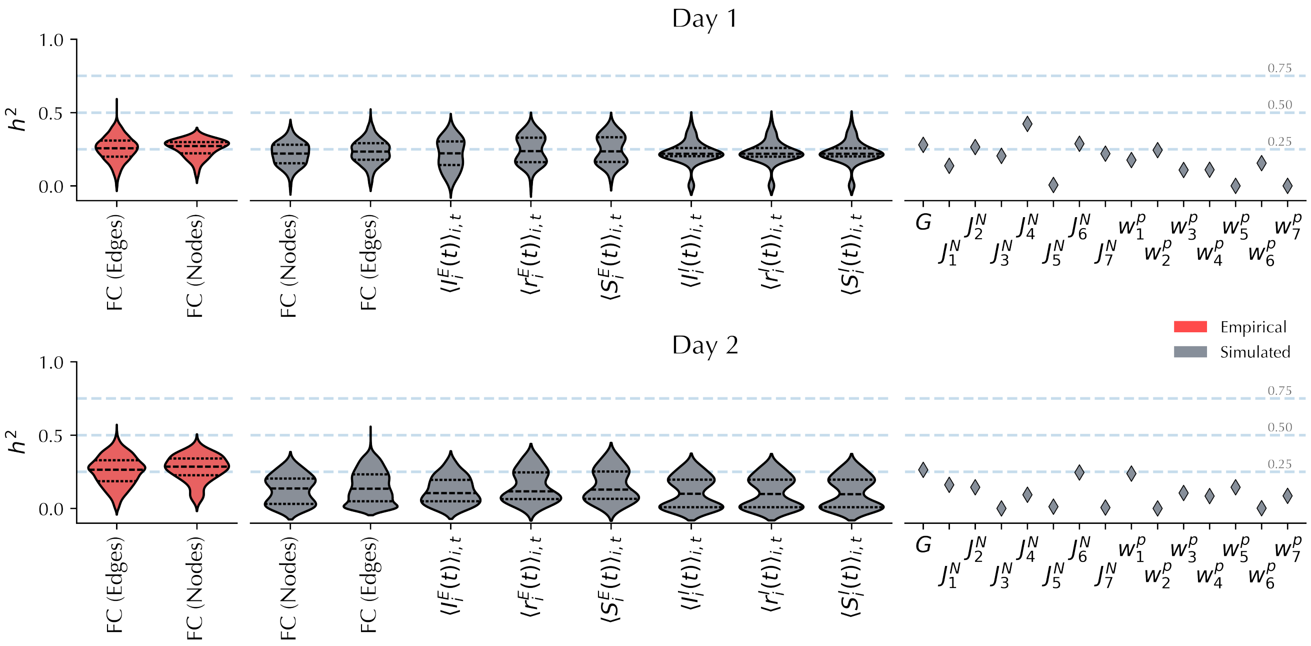


**Fig. S4. Session-specific heritability of empirical and simulated features.** Heritability estimates ($h^2$) are shown separately for data from day 1 (*top*; N = 402; age: 29.1±3.3; 58.2% female) and day 2 (*bottom*; N = 378; age: 29.2±3.4; 58.7% female). Empirical features are shown in red and simulated features in grey. Violin plots represent the distribution of values across connectome edges or network nodes, with black lines indicating medians and quartiles. Diamonds indicate point estimates.

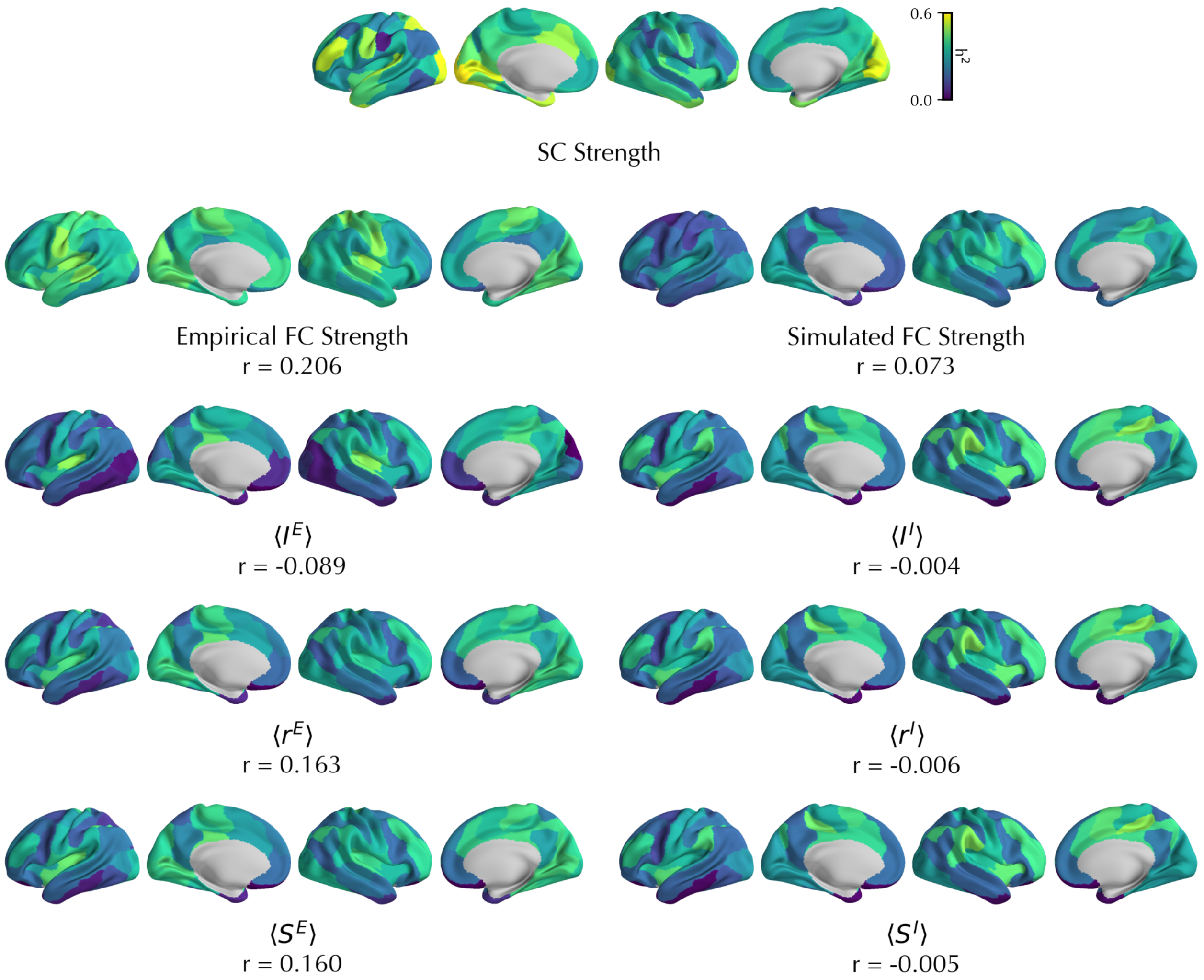


**Fig. S5. Spatial pattern of heritability measures.** *Top*: Regional heritability measure ($h^2$) of structural connectivity (SC) strength. *Bottom*: Regional heritability of empirical and simulated measures based on data averaged from day 1 and day 2 scans. Reported statistics below each map indicate Pearson correlations of regional heritability scores with the SC strength heritability map. No significant associations ($p_{spin,FDR} < 0.05$) were observed.

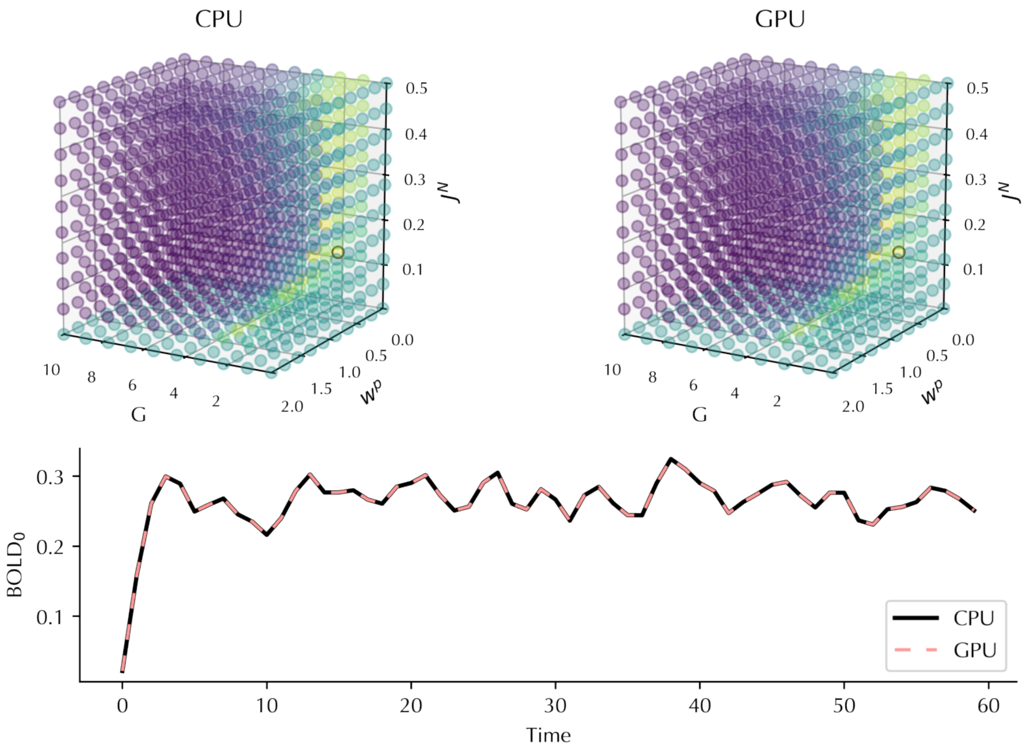


**Fig. S6. Identity of CPU and GPU outputs.** *Top*: Grids of negative cost calculated on CPU (*left*) and GPU (*right*) are shown. The grids were perfectly correlated (r = 1.0) and had a mean absolute difference of 7e-14. *Bottom*: BOLD time series in the first node of an example simulation calculated on CPU (solid black) and GPU (dashed light red) are shown and indicate their identity.
